# The catheterized bladder environment promotes Efg1- and Als1-dependent *Candida albicans* infection

**DOI:** 10.1101/2021.06.01.446547

**Authors:** Alyssa Ann La Bella, Marissa Jeme Andersen, Nicholas C. Gervais, Jonathan Jesus Molina, Alex Molesan, Peter V. Stuckey, Lauren Wensing, Clarissa J. Nobile, Rebecca S. Shapiro, Felipe Hiram Santiago-Tirado, Ana Lidia Flores-Mireles

## Abstract

Catheter-associated urinary tract infections (CAUTIs) account for 40% of all hospital-acquired infections. Given that 20-50% of all hospitalized patients receive a catheter, CAUTIs are one of the most common hospital-acquired infections and a significant medical complication as they result in increased morbidity, mortality, and an estimated annual cost of $340-370 million. *Candida spp*. – specifically *Candida albicans* – are a major causative agent of CAUTIs (17.8%), making it the second most common CAUTI uropathogen. Despite this frequent occurrence, the cellular and molecular details of *C. albicans* infection in the CAUTI microenvironment are poorly understood. Here, we characterize fungal virulence mechanisms and fungal biofilm formation during CAUTI for the first time. We found that the catheterized bladder environment triggers *Candida* virulence programs and robust biofilm formation through Efg1-dependent hyphal morphogenesis and Als1, an Efg1-downstream effector. Additionally, we show that the adhesin Als1 is necessary for *in vitro* and *in vivo C. albicans* biofilm formation dependent on the presence of fibrinogen (Fg), a coagulation factor released in the bladder due to the mechanical damage caused by urinary catheterization. Furthermore, in the presence of Fg, overexpression of *ALS1* in *C. albicans* led to enhanced colonization and dissemination, while deletion of *ALS1* reduced both outcomes during CAUTIs. Our study ultimately unveils the mechanism that contributes to fungal CAUTI, which may provide more effective targets for future therapies to prevent these infections.

## INTRODUCTION

*Candida albicans* is one of few fungal species that can cause serious infection in immunocompromised individuals and immunocompetent people with implanted medical devices (*1*). Normally, *C. albicans* is part of the healthy human microbiota, asymptomatically colonizing the gastrointestinal (GI) tract, reproductive tract, oral cavity, and skin of most humans (*1–3*). However, dysbiosis in the host environment, such as alterations in host immunity, stress, or resident microbiota, promotes *C. albicans* colonization and overgrowth, leading to biofilm formation (*1*).

*C. albicans* produces highly structured biofilms composed of multiple cell morphologies: yeast, pseudohyphae, and hyphae (*4, 5*). Hyphal formation, hyphae-specific gene expression, and biofilm formation are hallmarks of virulence, invasion, and immune evasion, resulting in tissue damage and invasive infection (*6*). *C. albicans* biofilm formation is initiated by the adherence of yeast cells to a surface, forming microcolonies, and is followed by proliferation of hyphae and pseudohyphae (*6*). Hyphal growth is triggered via numerous environmental signals including serum, neutral pH, and CO_2_, and is controlled by master transcriptional regulators such as Enhanced filamentous growth protein 1 (Efg1) (*7*). Biofilm maturation occurs when the hyphal scaffold is encased in an extracellular polysaccharide matrix (*8*), and the final step involves dispersion of yeast cells to seed planktonic infections or to establish biofilms at new locations in the surrounding environment (*6*).

These robust and highly structured biofilms often develop on implanted devices, allowing *C. albicans* to colonize urinary and central venous catheters, pacemakers, mechanical heart valves, joint prostheses, contact lenses, and dentures (1-3). Biofilm formation on these devices predisposes patients to bloodstream infections and leads to invasive systemic infections of tissues and organs (*1*). *Candida* biofilms on devices pose a serious threat since they are more resistant to antifungal therapy and can evade protection from host defenses (*8*). Newer antifungal drug classes, such as the echinocandins, that target the β-1,3 glucan component of the cell wall have shown some efficacy against *C. albicans* biofilms (*2, 9*); however, echinocandin-resistant *Candida spp.* have been recently emerging (*10*).

The 2016 National Healthcare Safety Network review found that *Candida spp.,* specifically *C. albicans,* has become the second most prevalent uropathogen causing 17.8% of catheter associated urinary tract infections (CAUTIs) (*11*). CAUTIs are the most common nosocomial infection with more than 150 million individuals acquiring these infections per year (*12*). The use of catheters and subsequent infections are not limited to hospital settings. In nursing homes, 11.9% of residents use indwelling catheters (*13*), and ~50% of catheterized residents will experience symptomatic CAUTIs (*14*). Despite *Candida spp*. prevalence in CAUTIs, there are no studies available regarding the pathogenesis of *Candida spp*. during CAUTI, and the roles of host and fungal factors remains unclear (*11, 15*).

Urinary catheterization causes dysbiosis of the bladder environment by compromising the urothelium, inducing inflammation, interfering with normal micturition (voiding), and disrupting host defenses in the bladder (*16–18*). The inflammation response caused by catheterization exposes epithelial receptors and recruits host factors that can be recognized by the pathogen, enabling microbial colonization and persistence within the urinary tract (*16–18*). Specifically, host clotting factor 1, fibrinogen (Fg), is released into the bladder to heal damaged tissues and prevent bleeding due to catheter-induced inflammation in both mice and humans (*19, 20*). Due to the constant mechanical damage caused by the urinary catheter, Fg accumulates in the bladder and deposits onto the catheter, where it is exploited by uropathogens (*19–22*). In fact, in an *ex vivo* study, we showed that a *C. albicans* CAUTI isolate colocalizes with Fg on urinary catheters (*20*). This suggests that aspects of the catheterized bladder environment induce changes in *C. albicans* virulence programming, prompting us to study potential fungal factors for CAUTI pathogenesis.

Here, we characterized the growth, morphology, and biofilm formation in urine, as well as the ability to cause CAUTIs, of five *C. albicans* clinical and laboratory strains. We found that urine supported *C. albicans* growth, activated virulence by inducing hyphal morphology, and enhanced Fg-dependent biofilm formation mediated by the transcriptional regulator Efg1. Furthermore, we found that catheterized-induced inflammation, specifically Fg, is necessary for fungal colonization of the urinary tract and further demonstrated that Efg1-dependent hyphal formation is critical for CAUTI establishment. Importantly, we identified that the Efg1-induced adhesin, Als1, is important for mediating Fg-fungal interactions *in vitro* and *in vivo* and its expression is crucial for fungal CAUTIs. This suggests that the development of therapies that target the Als1-mediated fungal-Fg interactions could potentially be useful in the treatment of *C. albicans* CAUTIs.

## RESULTS

### *C. albicans* survival and growth in urine promotes hyphal formation

*C. albicans* can infect a wide range of regions within the human body including the oropharyngeal area, gastrointestinal tract, intra-abdominal area, skin, genitals, and urinary tract, demonstrating its plasticity to survive and replicate in different host environments (*1, 2, 7, 12, 23, 24*). Since *C. albicans* is a prominent CAUTI pathogen, we compared growth and morphology in nutrient-rich and restrictive urine conditions using three urinary clinical isolates (Pt62, Pt65, and PCNL1 (*20, 25*)) and two laboratory strains (DAY286 and SC5314 (**Table S1**)). Since hyphal formation is important for promoting disease, we used a hyphal defective double deletion mutant strain in the SC5314 background, *efg1*Δ/Δ*cph1*Δ/Δ, to test whether hyphal formation is important in the catheterized bladder environment (**Table S1**). Additionally, microenvironment conditions induce *C. albicans* morphological changes that are associated with virulence (*1, 5*). *C. albicans* can exhibit different morphologies (*6*), including yeast, pseudohyphae, and hyphae (*5*). The hyphal morphology is associated with invasive growth and increased fungal virulence (*7, 26*). Therefore, we also determined how the bladder environment affects *C. albicans’* morphology.

For growth analysis, yeast extract peptone dextrose (YPD), a standard *C. albicans* growth medium, and BHI, the isolation medium for the clinical strains, were considered rich environments under both static and shaking conditions. Static growth was used to mimic the bladder environment and shaking growth was used as a comparison with standard lab culture conditions. For restrictive environments, we used human urine grown statically, supplemented with 10% human serum, amino acids (AA), bovine serum albumin (BSA), or Fg to mimic plasma protein extravasation in the catheterized bladder (*21*). Serum albumin and Fg were chosen because they are two of the most abundant host proteins on catheters retrieved from humans and mice, and it has been shown that other uropathogens use them as nutrient sources (*19, 21, 27, 28*). Lastly, we used AA as a general nitrogen source. Samples were taken at 0, 24, and 48 hours to assess growth by enumeration of colony forming units (CFUs) and morphology by fluorescent microscopy. As expected, *C. albicans* strains grew in higher densities in rich media with aeration while growth in static rich media was similar to any urine condition (**Fig. 1 and S1**). Growth in restrictive environments varied widely, but all strains were able to grow to some extent. Human serum promoted growth of all strains by 24 hrs and of most strains by 48 hrs. A subsequent decline was observed by 48 hrs for Pt65 and SC5314, possibly because all nutrients were consumed rapidly (**Fig. 1A-F**). BSA and AA moderately promoted growth in all strains except for DAY286 where growth was inhibited when compared to growth in urine alone (**Fig. 1A, D-F**). Fg enhanced growth of all strains compared to urine alone (**Fig. 1A-F**).

**Figure 1.**
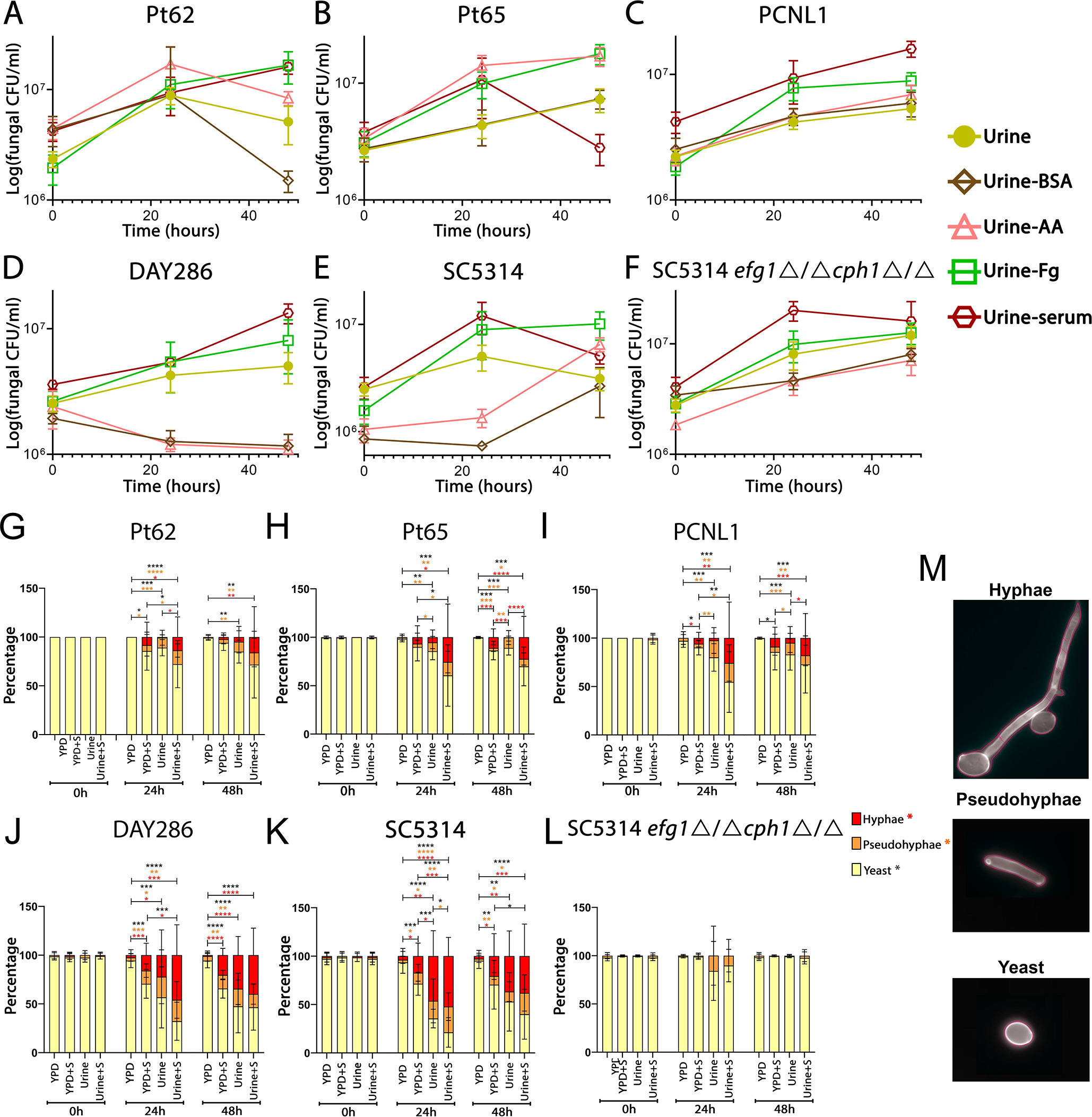
*C. albicans* grows, survives, and transitions to the hyphal morphology when grown in urine. **(A-F)** Growth curves of *C. albicans* strains Pt62 (A), Pt65 (B), PCNL1 (C), DAY230 (D), SC5314 (E), and SC5314 *efg1*Δ/Δ*cph1*Δ/Δ (F), grown in human urine conditions alone, or urine conditions supplemented with 10% human serum (serum), fibrinogen (Fg), bovine serum albumin (BSA), or amino acids (AA). Fungal growth was determined by CFUs enumeration after 0, 24, and 48 hours. Except when indicated, all strains were grown under static conditions. Data presented shows the mean and standard error of the mean derived from three independent experiments with at least three technical replicates. **(G-M)** The morphology of *C. albicans* strains were evaluated after 0, 24, and 48 hours of growth in urine and YPD with or without 3% human serum. Images (consisting of a 3 × 3 tiled region, i.e. 9 fields of view) were randomly acquired and at least three images were analyzed per condition. The total number of cells per phenotype were summed and divided by the total number of cells to give the overall percentage of yeast, pseudohyphae, and hyphae (**M**). Differences between groups were tested for significance using the Mann-Whitney U test. *, P < 0.05; **, P < 0.005; ***, P < 0.0005; and ****, P < 0.0001. P-value colors represent the comparison between the different populations: hyphae (red), pseudohyphae (orange), and yeast (black).

For morphology analysis, YPD and urine, with or without serum, were used as rich and restrictive environments, respectively. The YPD conditions served as negative and positive controls for filamentation with the absence or presence of serum, respectively (*29–31*). Cell morphology analysis was done by quantifying the percentages of yeast, pseudohyphal, or hyphal forms (**Fig. 1G-L, Table S2**). All strains showed predominantly yeast morphologies in YPD medium, and YPD with serum induced pseudohyphal and hyphal morphologies in all strains, except Pt65 and SC5314 *efg1*Δ/Δ*cph1*Δ/Δ. Notably, our analysis showed that urine conditions promoted pseudohyphal and hyphal formation in all strains (**Fig.1G-K, Table S2**) except SC5314 *efg1*Δ/Δ*cph1*Δ/Δ (**Fig. 1L, Table S2**). Pseudohyphal and hyphal morphologies were further induced when urine was supplemented with human serum in all WT lab and clinical strains. This suggests that the catheterized environment triggers *C. albicans* to transition from the yeast to hyphal morphology.

### Fibrinogen enhances *C. albicans* biofilm formation

During candidiasis, *C. albicans* pseudohyphal and hyphal formation, observed on catheters from CAUTI patients (*15, 25*) and rats (*32, 33*), induces expression of virulence genes including adhesion factors (*4, 6, 34, 35*). Furthermore, our *ex vivo* study showed that the *C. albicans* PCNL1 clinical isolate colocalized with Fg, a known binding and biofilm platform for diverse uropathogens (*19, 20, 27, 36, 37*). Together, this suggests that Fg may be an important factor promoting *C. albicans* CAUTI pathogenesis. Thus, we hypothesized that urine conditions induce factors responsible for Fg-binding and biofilm formation.

Biofilm formation was compared in rich (YPD and BHI) and restrictive (human urine) media as well as between BSA- and Fg-coated microplates (**Fig. 2A-F**). At 48 hrs, immunostaining was performed to assess fungal biofilm biomass (*38*). We found that Fg promoted biofilm formation of most strains under all conditions and was further enhanced by human urine (**Fig. 2A-C**). Fg enhanced biofilm formation in DAY286 in YPD and urine but not in BHI **(Fig. 2D)**. SC5314 showed enhanced biofilm formation with Fg in all conditions **(Fig. 2E)**. Importantly, SC5314 *efg1*Δ/Δ*cph1*Δ/Δ formed enhanced biofilms in rich media when coated with Fg but was unable to do so in urine, regardless of the coated surface (**Fig. 2F**). This suggests that not only is filamentation important for biofilm formation in urine environments, but that biofilm formation mechanisms differ between urine and rich environments.

**Figure 2.**
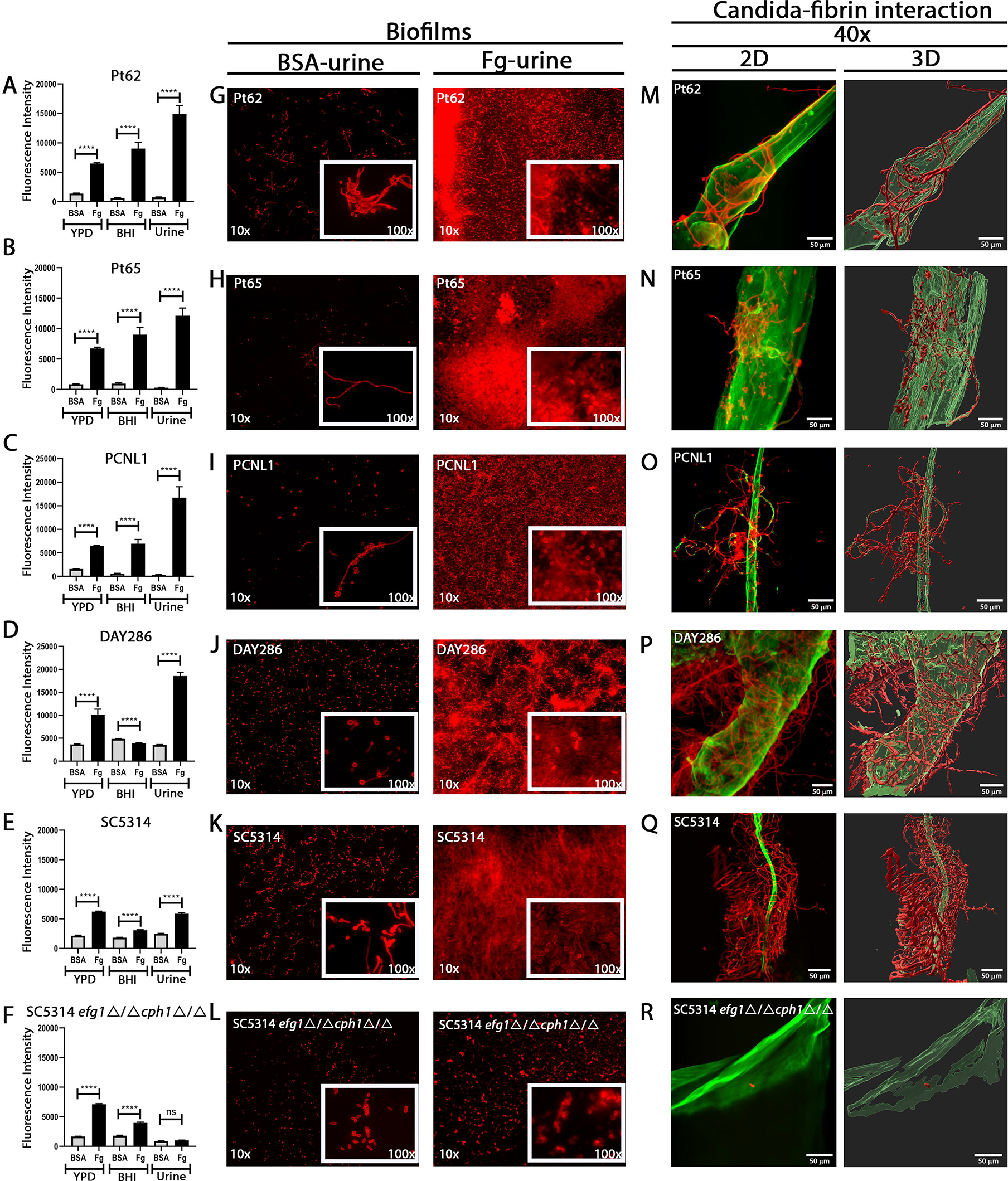
Fibrinogen/fibrin interaction with *C. albicans* enhances biofilm formation. (**A-F**) Immunostaining analysis of biofilm formation on BSA- or Fg-coated microplates by *C. albicans* strains when grown in YPD, BHI, or human urine. At 48 hrs, *C. albicans* biofilm formation was measured via fluorescence intensity using anti-*Candida* antibodies. Data presented shows the mean and standard error derived from three independent experiments with 24 technical replicates. Differences between groups were tested for significance using the Mann-Whitney U test. ****, P<0.0001; ns, not statistically different. (**G-L**) Microscopic visualization of 48 hrs *C. albicans* biofilms biomass on BSA- or Fg-coated glass bottom petri dishes grown in urine using anti-*Candida* antibodies. White squares represent the zoom-in area used for the higher magnification (x). (**M-R**) Microscopic visualization and 3D reconstruction of 48 hrs *C. albicans* biofilms on fibrin fibers/meshes grown in human urine using antibodies against Fg (anti-Fg; green) and *C. albicans* (anti-*Candida*; red). Scale bars: 50 μm.

Further analysis of biofilm formation was performed on *C. albicans* biofilms grown in urine on plates coated with either BSA, Fg, or fibrin meshes. Fibrin meshes were added because in damaged tissue environments like the catheterized bladder, Fg is converted into fibrin fibers or meshes to stop bleeding and allow tissue healing (*39*). Visual examination of biofilm formation on BSA-, Fg- or fibrin-coated plates showed that Fg- and fibrin-coated plates supported robust biofilm formation composed of yeast, pseudohyphae, and hyphae, while on BSA-coated plates, sparse monolayers or small aggregates of cells were observed (**Fig. 2G-K, M-Q; Fig. S2**). However, SC5314 *efg1*Δ/Δ*cph1*Δ/Δ showed a small yeast-exclusive monolayer of cells regardless of BSA, Fg, or fibrin presence (**Fig. 2L and R; Fig. S2)**. Together, this suggests filamentation is important for Fg- and fibrin-dependent biofilm formation and that the regulation of Fg-binding adhesins expressed during hyphal morphogenesis is controlled by Efg1, Cph1, or their downstream effectors.

### *C. albicans* hyphal formation and Fg interaction are critical for establishment of CAUTI

Based on these results and the importance of biofilm formation for infections such as CAUTIs, we hypothesized that the *C. albicans* hyphal-deficient mutant strain would be unable to colonize the bladder and catheter. To test this hypothesis, we compared the ability of the clinical and laboratory strains to colonize the bladder both in the presence and absence of a catheter using our established mouse model (*19, 21, 27, 28, 36, 37, 40–43*). Mouse bladders were challenged with one of the strains, and 24 hours post infection (hpi), the mice were euthanized, and fungal colonization was quantified via CFUs. Our results showed that catheterization significantly promoted bladder colonization by all the clinical and lab strains compared to the bladder colonization in non-catheterized mice (**Fig. 3**). Importantly, colonization by the hyphal-deficient mutant strain, SC5314 *efg1*Δ/Δ*cph1*Δ/Δ, was significantly impaired (**Fig. 3E**), and in the absence of a catheter, behaved and colonized to the same extent as the WT strain. Since our mouse model of CAUTI allows us to assess dissemination, we analyzed the fungal burden of the kidneys, spleen, and heart after 24 hpi (**Fig. 3**). We found that urinary catheterization significantly contributed to fungal spread of DAY286 to the kidneys and spleens (**Fig. 3D**) and spread of SC5314 to the kidneys (**Fig. 3E**). Kidney colonization by Pt62 and Pt65 was 2-3 logs higher than non-catheterized mice, trending to significance. Importantly, the hyphal-deficient mutant strain, SC5314 *efg1*Δ/Δ*cph1*Δ/Δ, did not show differential dissemination between catheterized and non-catheterized mice (**Fig. 3E**).

**Figure 3.**
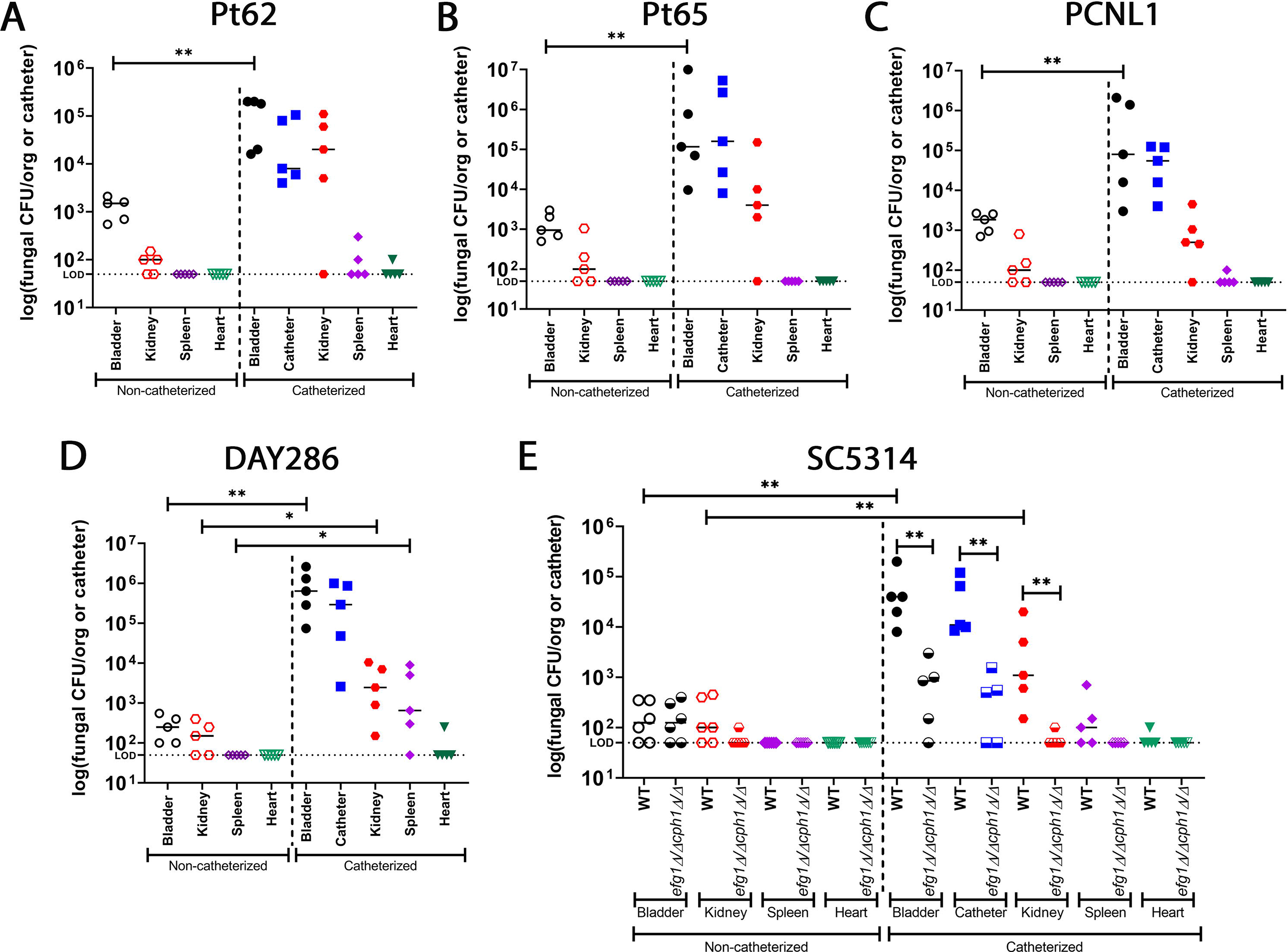
Urinary catheterization promotes fungal colonization and hyphal formation is required for CAUTI. (**A-D**) Fungal burden of clinical and lab strains on the harvested organs and catheter. (**E**) Fungal burden comparison between SC5314 WT and SC5314 *efg1*Δ/Δ*cph1*Δ/Δ strains. Values represent means ± SEM. The Mann-Whitney U test was used; *, P < 0.05 was considered statistically significant. **, P < 0.005; ns, values were not statistically significantly different. The horizontal bar represents the median value. The horizontal broken line represents the limit of detection of viable fungi. LOD; limit of detection. Animals that lost the catheter were not included in this work.

To further understand *C. albicans* morphogenesis, interaction with Fg, and spatial colonization in the bladder during CAUTI, we performed histological analyses and immunofluorescence (IF) microscopy of catheterized bladders at 24 hpi. Bladder colonization was exceptionally robust and was visible in the H&E-stained whole bladders (blue arrow heads) (**Fig. 4**). Consistently, our IF analysis showed the presence of *C. albicans* hyphal and pseudohyphal morphologies in the lumen of the bladder in all clinical and laboratory strains (**Fig. 4A-E**), except the hyphal-deficient mutant strain, SC5314 *efg1*Δ/Δ*cph1*Δ/Δ (**Fig. 4F**). Furthermore, *C. albicans* cells in the catheterized bladder were found associated with Fg (**Fig. 4**, see **Fig. S3-S8** for zoomed-in images). These data indicate that the changes induced by catheterization promote fungal colonization of the urinary tract and further demonstrate that hyphal morphogenesis, as well as Efg1 and Cph1 regulated pathways, are crucial for CAUTI establishment.

**Figure 4.**
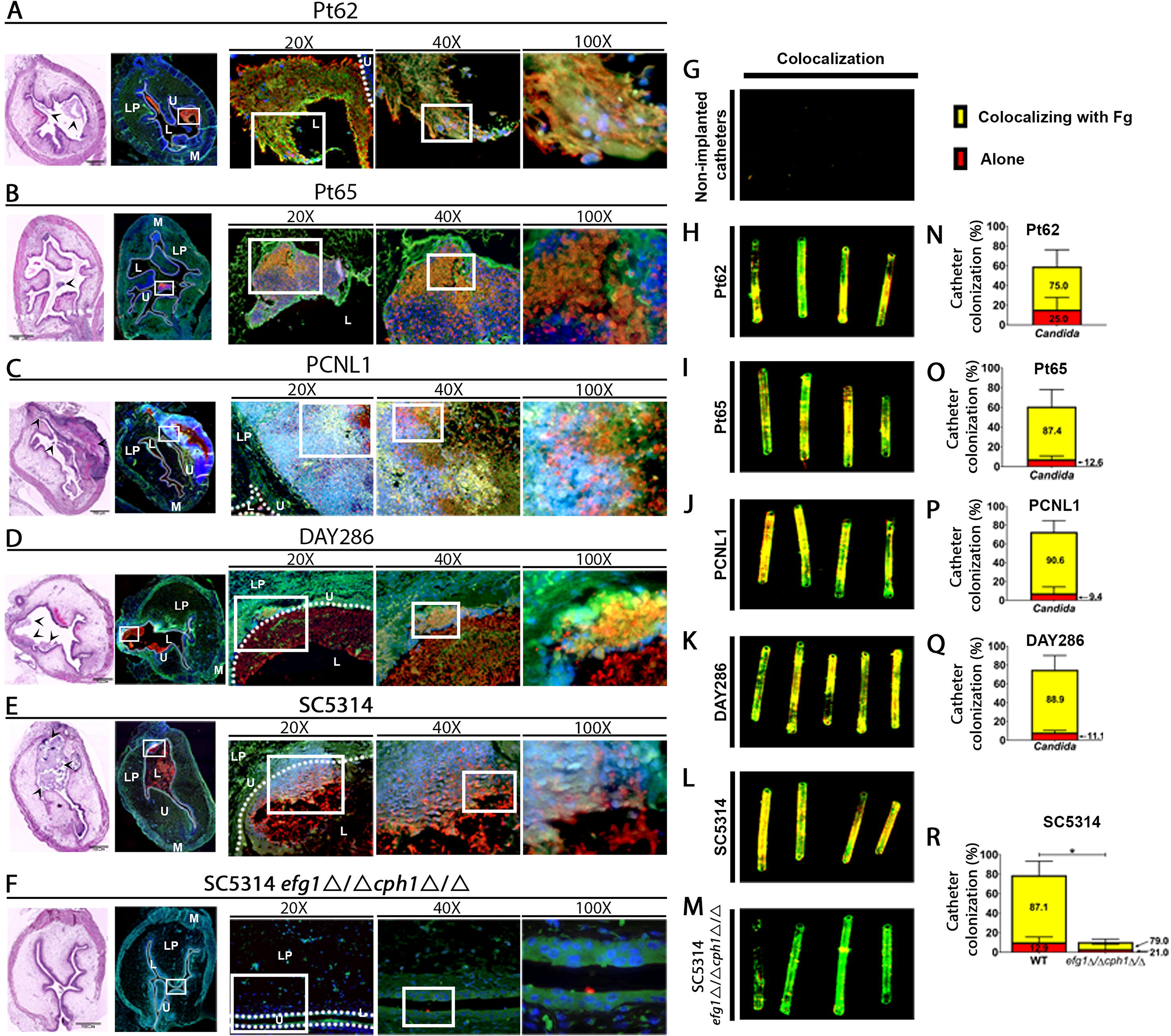
Hyphal *C. albicans* cells invade the lumen of the catheterized bladder and interact with fibrinogen on the bladders and catheters. (**A-F**) Implanted and infected bladders and catheters were recovered at 24 hpi. Bladders were subjected to analysis by H&E and IF staining. For the IF analysis antibody staining was used to detect Fg (anti-Fg; green), *C. albicans* (anti– *Candida*; red), and neutrophils (anti-Ly6G; white). Staining with DAPI (blue) delineated the urothelium and cell nuclei (representative images). The white broken line separates the bladder lumen (L) from the urothelium surface (U), the lamina propria (LP), and muscularis (M). H&E stained bladder scale bars, 700 μm. White squares represent zoomed-in areas of higher magnification (20x, 40x, 100x). Blue arrow heads indicate *C. albicans* colonization. (**G-M**) Implanted catheters were stained with antibodies to detect Fg (anti-Fg; green) and *C. albicans* (anti-Candida; red). (**N-R**) Quantification of fungal colocalization with deposited Fg on the catheter. The Mann-Whitney U test was used to analyze catheter colonization between SC5314 WT and hyphal mutant; *, P < 0.05; Values represent the means ± standard deviation derived from co-localization of the catheter segments. Non-implanted catheters were used as negative controls (**G**).

### *C. albicans* interacts with Fg on the catheter surface *in vivo* during CAUTI

Based on our *in vitro* Fg-binding results (**Fig. 2**), we assessed *C. albicans*-Fg interactions *in vivo* on the catheter during CAUTI. Catheters from mice infected with each strain were retrieved 24 hpi, stained, and imaged. Except for the hyphal-deficient mutant strain, we found that all strains formed a robust biofilm on the implanted catheter (**Fig. 4G-R** merge images; **Fig. S9**, single channels), colonizing 59% to 79% of the surface of the catheter (**Table S3**). SC5314 WT showed 78.9 ± 14% catheter colonization, while SC5314 *efg1*Δ/Δ*cph1*Δ/Δ showed 10.4 ± 7.5% catheter colonization, exhibiting a significant defect in colonization (**Fig. 4L-M, R; Table S3**). Our IF analysis showed that *C. albicans* strains preferentially bound to deposited Fg on the catheters (**Fig. 4H-M and Fig. S9**). Quantification of the staining found that 75-91% of the *C. albicans* strains were colocalized with deposited Fg (**Fig. 4N-R and Fig. S9**). Although the hyphal-deficient strain was only able to colonize 10% of the catheter (**Fig. 4M, Table S3**), its cells were colocalizing with Fg (**Fig. 4R, Table S3**). This result further corroborates the importance of hyphal morphogenesis for biofilm formation and Fg for catheter colonization during CAUTI.

### Efg1 is critical for filamentation and for Fg-dependent biofilm formation in urine, and in the development of CAUTI

We next explored the independent contributions of Efg1 and Cph1 to hyphal morphogenesis, Fg-dependent biofilm formation, and colonization of the catheterized bladder environment (**Fig. 5 and S10**). Using single deletion strains with their corresponding WT, we found that the *cph1*Δ/Δ strain produced pseudohyphae and hyphae similar to WT while the *efg1*Δ/Δ strain was predominantly in the yeast-form throughout all conditions (**Fig. S10)**. Fg-dependent biofilm formation was assessed and the *cph1*Δ/Δ strain exhibited a ~40% decrease in biofilm formation when compared to the WT strains while the *efg1*Δ/Δ strain showed a ~95% reduction, and the *EFG1* complemented strain restored biofilm formation to WT levels (**Fig. 5A**). Furthermore, analysis of fibrin interactions showed that the *cph1*Δ/Δ strain produced a robust hyphal colonization surrounding the Fg fibers, differing from the *efg1*Δ/Δ strain, which showed few yeast cells bound directly to the Fg mesh (**Fig. 5B and Fig. S2**). Analysis of colonization in our mouse model showed that the *efg1*Δ/Δ strain significantly impaired bladder and catheter colonization while the *EFG1* complemented and *cph1*Δ/Δ strains displayed comparable colonization to the WT strain (**Fig. 5C**). Based on the role of Efg1 in CAUTI, *EFG1* expression levels were analyzed in the 24 hpi catheterized and infected bladders with WT, *efg1*Δ/Δ, and *EFG1* complemented strains. We found that *EFG1* is expressed during bladder infection with the WT and *EFG1* complemented strains and that *EFG1* transcripts were undetectable (ND) in the *efg1*Δ/Δ infected bladders as expected (**Fig. 5E**). Furthermore, analysis of bladder tissue showed that the *cph1*Δ/Δ strain infected bladders had abundant hyphal burdens similar to the WT strain while the *efg1*Δ/Δ strain infected bladders showed defective colonization by sporadic yeast cells (**Fig. 5F, S11-S13**). These phenotypes exhibited by the *efg1*Δ/Δ strain mimicked the phenotypes shown by the *efg1*Δ/Δ*cph1*Δ/Δ strain, indicating that Efg1 is the major transcriptional regulator responsible for the yeast to hyphal transition and for Fg-dependent biofilm formation in the catheterized bladder environment.

**Figure 5.**
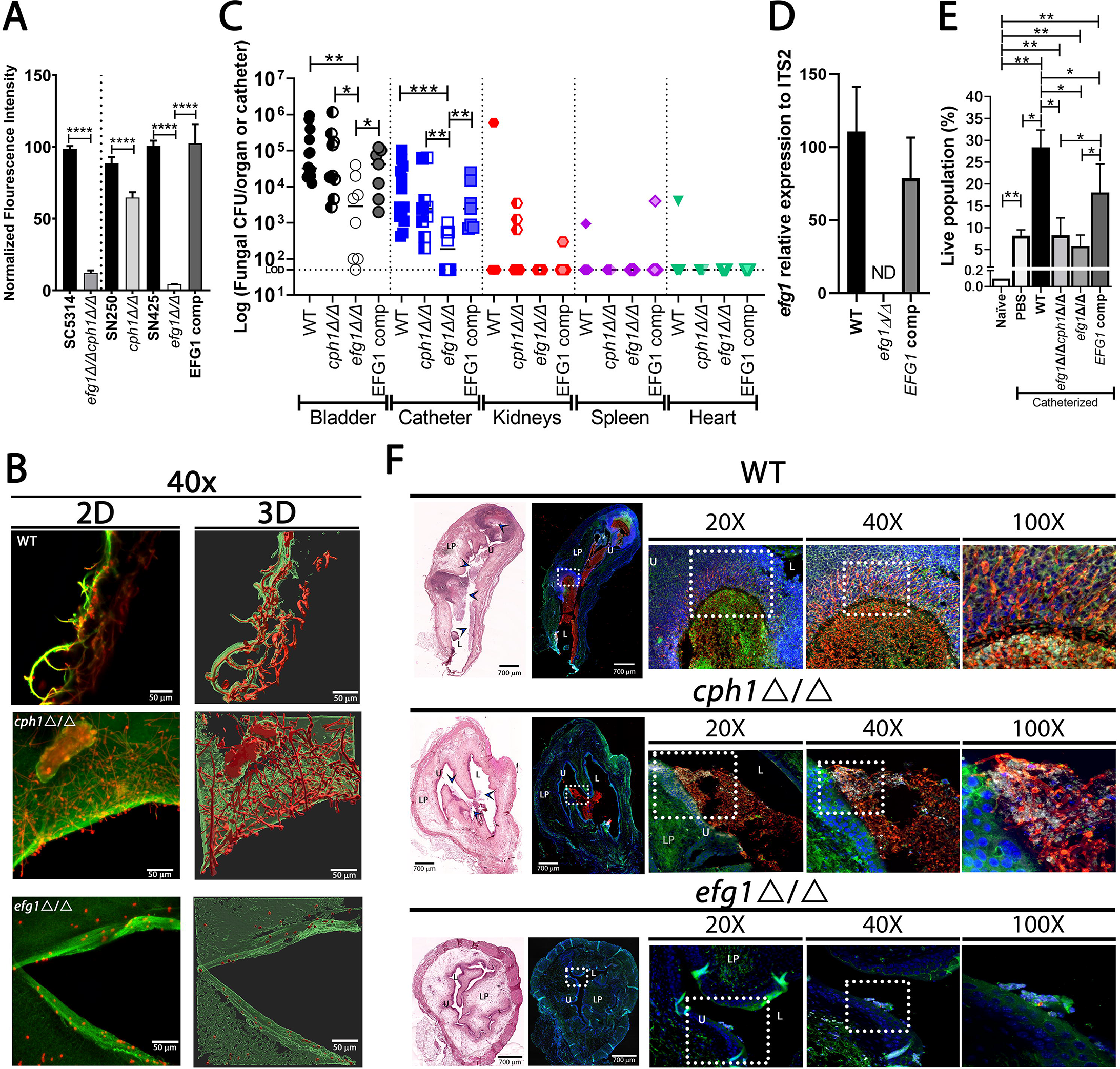
Efg1 is responsible for inducing hyphal morphogenesis and is critical for establishment of CAUTI. (**A**) Comparison of biofilm formation on Fg-coated microplates by *C. albicans* strains when grown in urine for 48 hours. (**B**) Microscopic visualization and 3D reconstruction of *C. albicans-*fibrin/meshes interaction grown in urine using antibodies against Fg (anti-Fg; green) and *C. albicans* (anti-*Candida*; red). Scale bars: 50 μm for 40x. (**C**) Mice were catheterized and challenged with 1×10^6^ CFUs of each strain, and burdens in organs or catheters were quantitated as the number of CFUs recovered. **(D)** RNA was extracted from 24 hpi catheterized and infected bladders with either the WT, *efg1*Δ/Δ, and *EFG1* complemented strains and *EFG1* expression analysis was assessed by qRT-PCR. **(E)**Neutrophil recruitment to the bladder in response of catheterization and infection. (**F**) Mouse bladders were harvested following infection and catheter implantation. Bladders were subjected to analysis by H&E and IF staining (anti-Fg, green; anti-*Candida*, red; anti-Ly6G, white; DAPI, blue). Values represent means ± SEM. The Mann-Whitney U test was used; P < 0.05 was considered statistically significant. *, P < 0.05; **, P < 0.005; ***, P < 0.0005. The horizontal bar represents the median value. The horizontal broken line represents the limit of detection of viable fungi. Animals that lost the catheter were not included in this work. ND, Not detectable.

Neutrophils are highly recruited into the catheterized bladder (*21, 28, 44*) and neutropenic patients are more susceptible to *C. albicans* and bacterial dissemination originating from CAUTI (*45–48*). We next analyzed neutrophil recruitment into the catheterized and infected bladder via IF and flow cytometry (FC). Imaging of bladders showed that neutrophils were highly recruited into the bladder, specifically in the areas with fungal colonization (**Fig. 4** and **Fig. S3-S8**). We also observed *C. albicans* cells breaching the urothelium where they encounter a strong neutrophil response (**Fig. 4C-E** and **Fig. S5-S7**). For example, PCNL1 was able to breach the bladder lamina propria, inducing massive neutrophil recruitment to contain the infection (**Fig. 4C** and **Fig. S5**). Pt62 and Pt65 strains were primarily found in the bladder lumen and their cells were interacting with Fg and neutrophils (**Fig. 4A-B** and **Fig. S5-S6**). On the other hand, robust fungal colonization and neutrophil recruitment was not observed in the *efg1*Δ/Δ*cph1*Δ/Δ strain or *efg1*Δ/Δ strain infected bladders (**Fig. 4D, 5F, S8**, and **S13**). Based on this data, using flow cytometry, we quantified and compared bladder neutrophil recruitment in naïve and catheterized bladders that were PBS-mock infected or infected with the WT, *efg1*Δ/Δ*cph1*Δ/Δ, *efg1*Δ/Δ, and *EFG1* complemented strains (**Fig. S14**). We observed that the catheterized bladder by itself (PBS-mock infected) induced neutrophil recruitment; however, the infected bladder with the WT strain further increased neutrophil recruitment (**Fig. 5E**), as we have observed via IF (**Fig. 5F**). Furthermore, neutrophil recruitment in the bladder infected with the *efg1*Δ/Δ*cph1*Δ/Δ and *efg1*Δ/Δ strains showed levels comparable to the PBS-mock infected condition, while the *EFG1* complemented strain was not significantly different from the WT strain (**Fig. 5D**). These data demonstrate that in the catheterized bladder, infection with *C. albicans* leads to the recruitment of neutrophils to the site of infection, and the virulent hyphal and pseudohyphal morphologies are driven by Efg1.

### An Efg1-regulated adhesin is critical for biofilm formation and persistence during CAUTI

We have demonstrated that *C. albicans* interacts with Fg and its presence enhances colonization in the catheterized bladder (**Fig. 2–3**), suggesting that urine conditions induce the expression of Fg-binding adhesins. To identify adhesins important for attachment, colonization, and persistence during CAUTI, we screened 12 adhesin mutant strains from the Shapiro library (*49, 50*) (**Fig. 6A**) and 27 adhesin and biofilm-related mutants from the Noble library (*51*) (**Fig. S15A**) for Fg-dependent biofilm formation under urine conditions. We found that of all adhesin mutant strains, the *als1*Δ/Δ strain exhibited the most severe biofilm formation defect under urine conditions (**Fig. 6A**), similar to the biofilm defects of the *efg1*Δ/Δ and *efg1*Δ/Δ*cph1*Δ/Δ strains, without compromising hyphal morphology (**Fig. 6A, 2F and Fig. S16**). Furthermore, the *ALS1* complemented strain rescued biofilm formation to WT levels (**Fig.6B**; *ALS1 comp*). In addition, we assessed biofilm formation by an *ALS1* overexpression strain (*ALS1* OE; obtained from the Shapiro lab (*52*)). This strain not only rescued but enhanced biofilm formation to 8-fold higher than the WT strain **(Fig. 6B)**. Increased biomass was clearly visible in the *ALS1* OE strain and due to insufficient antibody penetration, crystal violet was used to assess biomass (**Fig. S15B)**. Visualization of the WT, *als1*Δ/Δ, and *ALS1* OE strains interactions with fibrin showed that the *als1*Δ/Δ strain displayed less colonization of the fibrin meshes while the *ALS1* OE strain exhibited a more robust colonization than the WT strain (**Fig. 6C**).

**Figure 6.**
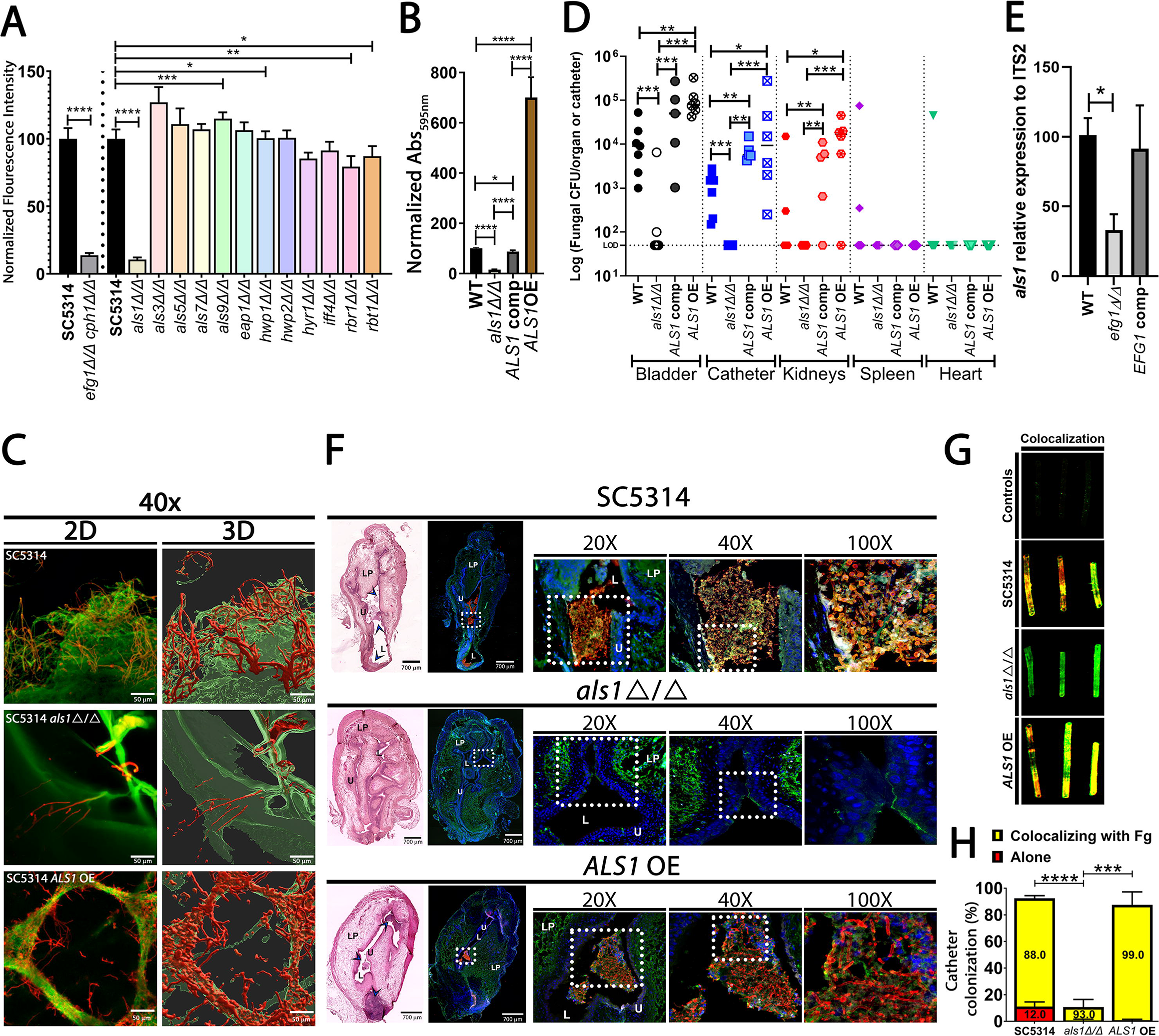
Als1 is critical for Fg-dependent biofilm formation and establishment of CAUTI. Biofilm formation analysis on Fg-coated microplates by *C. albicans* adhesin mutant strains (**A**) or the *ALS1* OE strain (**B**) when grown in human urine. At 48 hrs, *C. albicans* biofilm formation was measured via fluorescence intensity using anti-*Candida* antibodies (**A**) or crystal violet (**B**). (**A-B**) Data presented shows the mean and standard error derived from three independent experiments with 24 technical replicates. (**C**) Microscopic visualization and 3D reconstruction of *C. albicans-*fibrin/meshes interaction grown in urine using antibodies against Fg (green) and *C. albicans* (red). Scale bars: 50 μm for 40x. (**D**) Fungal burden in organs or catheters at 24 hpi were quantitated as the number of CFUs recovered. € RNA was extracted from 24 hpi catheterized and infected bladders with either the WT, *efg1*Δ/Δ, and *EFG1* complemented strains and *ALS1* expression analysis was assessed by qRT-PCR. (**F**) H&E and IF analysis of catheterized and infected bladders. IF staining was performed to detect Fg (green), *C. albicans* (red), and neutrophils (white). DAPI (blue) was used to stain the cell nuclei and delineated the urothelium (representative images). L, represents; U, urothelium; LP, the lamina propria; and M, muscularis. Bladder scale bars, 700 μm. White squares represent zoomed-in areas of higher magnification (20x, 40x, and 100x). Blue arrow heads on H&E indicate *C. albicans* bladder colonization. (**G**) *C. albicans* and Fg colocalization in implanted catheters using antibodies against Fg (anti-Fg; green) and *C. albicans* (anti-*Candida*; red). (**H**) Quantification of fungal colocalization with deposited Fg on the catheter. For biofilm formation, fungal burden and colocalization analysis values represent means ± SEM. The Mann-Whitney U test was used; *, P < 0.05; **, P < 0.005; ***, P < 0.0005; and ****, P < 0.0001. The horizontal bar represents the median value. (**D**) The horizontal broken line represents the limit of detection of viable fungi. Animals that lost the catheter were not included in this work.

We further compared the ability of the WT, *als1*Δ/Δ, *ALS1* complemented, and *ALS1* OE strains to cause CAUTI. The *als1*Δ/Δ strain displayed a significant defect in bladder and catheter colonization while the colonization of the *ALS1* complemented strain was similar to that of the WT; in addition, the *ALS1* OE strain showed a significant increase in bladder, catheter, and kidney colonization when compared to the WT strain (**Fig. 6D**). Als1 has been reported to be an Efg1-downstream target (58). To confirm this, *ALS1* expression levels were analyzed in 24 hpi catheterized and infected bladders with either WT, *efg1*Δ/Δ, and *EFG1* complemented strains. We found that *ALS1* expression significantly decreased in bladders infected with the *efg1*Δ/Δ strain compared with bladders infected with the WT and *EFG1* complemented strains (**Fig. 6E**). Furthermore, imaging of the *als1*Δ/Δ strain infected bladders did not show fungal colonization (**Fig. 6F**), whereas the *ALS1* OE strain infected bladders had abundant hyphal burdens similar to the WT strain (**Fig. 6F; Fig. S17-S19**). Imaging of catheters found that the WT strain colonization covered ~92.4% of the surface, the *als1*Δ/Δ strain covered ~10.73%, and the *ALS1* OE strain covered ~87.5%. Of that colonization, the fungal population colocalizing with deposited Fg was ~88.0% for the WT strain, ~93.0 for the *als1*Δ/Δ strain, and ~99.0% for the *ALS1* OE strain (**Fig. 6G-H and Fig. S20**). The residual binding to Fg in the *als1*Δ/Δ strain was similar to that of the *efg1*Δ/Δ*cph1*Δ/Δ strain, suggesting that this binding is mediated by other adhesins. Together, these data show that Als1 is induced by Efg1 and that Als1 is the Fg-binding protein critical for establishing colonization in the catheterized bladder and that modulation of Als1 expression affects the outcome of infection.

## DISCUSSION

### Candida spp

CAUTIs are increasing, most commonly in prolonged urinary catheterization and ICU settings (*53, 54*). Our results have shown that urinary catheterization significantly promotes colonization of the bladder by all strains to ≥10^5^ CFUs/ml, meeting the CDC’s standards of UTI criteria (*55*), whereas in the absence of the catheter, colonization by all strains was significantly lower at ≤10^3^ CFUs/ml (**Fig. 3**). It is possible that colonization by the WT lab strain and the clinical strains is even higher than the CFU enumeration indicates since hyphal cells are sticky multinucleated cells. Our model indicates that the prevalence of *C. albicans* in the infected bladder is due to increased colonization resulting from changes in the bladder environment caused by urinary catheterization.

In this study, we also found that Efg1-dependent hyphal morphogenesis and Als1-mediated interactions with Fg and fibrin are critical for fungal CAUTI establishment (**Fig. 5–6**). Importantly, Fg enhances *C. albicans* biofilm formation via interactions with the adhesin Als1. Additionally, urine promotes Efg1 hyphal morphogenesis which is essential for biofilm formation and infection during CAUTI. *C. albicans* hyphal formation induces strong neutrophil recruitment in the bladder to control the fungal infection. Thus, the catheterized bladder creates the ideal environment for *C. albicans* to colonize and persist in the host.

Host proteins have been shown to contribute to fungal biofilm formation (*33*). We demonstrated that Fg is accumulated in the bladder and deposited on the catheters (*11, 20*) and that the *C. albicans* PCNL1 clinical isolate colocalized with Fg deposited on urinary catheters retrieved from a patient with preoperative negative urine culture (*20*). Moreover, a study from the Andes group showed that Fg, as well as other host proteins, were associated with *C. albicans* biofilms in urinary catheters retrieved from rats (*8*). Therefore, we focused on understanding the role of Fg in *C. albicans* CAUTIs. We found that hyphal formation was critical for Fg-dependent biofilm formation in urine conditions but not in YPD and BHI media, while the hyphal defective mutant strain was not able to form Fg-dependent biofilms in urine, but could in rich media (**Fig. 1 and S1**). This seemingly contradictory result is not necessarily surprising since we have observed that laboratory growth media does not fully recapitulate conditions found within the host. For example, several factors critical for bacterial biofilm formation in CAUTI are dispensable for *in vitro* biofilm formation when using conventional laboratory growth media (*19, 28*). This difference could be related to adhesins that are expressed in the yeast cells that may contribute to biofilm growth in laboratory conditions. Thus, using conditions that closely mimic the *in vivo* environment is important to identify physiologically relevant determinants for CAUTI colonization and survival.

To further mimic the bladder environment, we incorporated the presence of host proteins, BSA and Fg. These proteins are found in the catheterized bladder due to the inflammation response caused by urinary catheter mechanical damage. Indubitably, we found that Fg enhances fungal biofilm formation in urine conditions and during urinary catheterization, and that this is mediated by the adhesin, Als1 (**Fig. 2, 6**). *C. albicans* expresses numerous adhesion proteins, including the known Fg-binding protein Mp58 (Pra1 in SC5314); however, in urine conditions, Als1 is the primary adhesin of Fg. We found that modulation of Als1 affected the outcome of biofilm formation and infection, with the *C. albicans als1Δ/Δ* strain displaying defective Fg-dependent biofilm formation and *in vivo* colonization. Conversely, the *ALS1* complemented strain and the *ALS1* overexpression strain rescued Fg-dependent biofilm formation and promoted CAUTI pathogenesis (**Fig. 6**). Previous structural studies showed that the N-terminal domain of Als1 interacts with Fg (*56, 57*), binding to the Fg γ-chain via protein-protein interactions, similar to Clf adhesins in *S. aureus* (*57, 58*). Als1 belongs to the agglutinin-like sequence (Als) protein family of cell-surface glycoproteins, which are important for adhesion to host, bacteria, and abiotic surfaces (*56*). Interestingly, Als3 and Als9-2 (an allelic variant of Als9) have been shown to interact with Fg (*56, 57*); however, they are dispensable in Fg-dependent biofilm formation in urine (**Fig. 6A**). This may suggest that these factors are not expressed under urine conditions. Our results indicate that Als1 is a Efg1-downstream target in the catheterized bladder environment. Studies have shown that Efg1 regulates *ALS3* expression under serum conditions (*59*) and that Efg1 directly binds to the *ALS3* promoter during biofilm formation in Spider media (*60*). Efg1 has both overlapping and distinct targets under different environmental conditions (*59*). Therefore, future studies dissecting the Efg1 regulatory network in the catheterized bladder will be crucial to understanding *C. albicans’* cellular signaling and to better manage the outcome of infection.

It has been established that urinary catheterization causes mechanical damage to the bladder, inducing epithelial wounding and triggering robust inflammation (*21, 61*). Additionally, clinical studies have detected *C. albicans* hyphae and biofilms on indwelling urinary catheters retrieved from patients with candiduria (*15, 62, 63*), suggesting fungal pathogenic activity during urinary catheterization. Here, we demonstrated that urine induces hyphal morphogenesis *in vitro* and *in vivo* (**Fig. 1 and Table S2**) and that Efg1-dependent hyphal programming is critical for bladder and catheter colonization (**Fig. 5**). Recently, deletion of *EED1* in *C. albicans* was shown to result in inhibition of hyphal formation but not virulence; this was attributed to the overexuberant growth of the *eed1*Δ/Δ strain (*64*). Since we observed hyphal invasive colonization in the catheterized bladder (**Fig. 4C-E; 5E**), in future experiments, we will explore whether the *eed1*Δ/Δ strain is able to cause CAUTI, promote bladder tissue invasion, colonization, and induce a strong neutrophil response.

Furthermore, strong neutrophil recruitment during hyphal colonization (for the WT, *cph1*Δ/Δ, and *EFG1* complemented strains) in the catheterized bladder was observed whereas colonization with hyphal deficient strains (*efg1*Δ/Δ*cph1*Δ/Δ and *efg1*Δ/Δ) showed similar neutrophil levels as the PBS-mock infected bladders (**Fig. 5**). This observation is important since invasive fungi (in the hyphal form) evoke an active host immune and inflammatory response while commensal fungi (in the yeast form) promote immune tolerance and “peaceful coexistence” with the microbiota (*65, 66*). Our results strongly suggest that *C. albicans* has pathogenic programming triggered by the catheterized bladder environment, resulting in fungal CAUTI.

During CAUTI, neutrophils are heavily recruited to the bladder as a defense mechanism against invading pathogens and their numbers increase with extended catheterization (*21, 44*). In neutropenic mice, the bacterial burden increases 10-fold during CAUTI, suggesting that neutrophils are controlling and containing the infection in the bladder (*21*). Similarly, neutropenic patients developed candidemia from candiduria, suggesting that bladder recruited neutrophils are critical to controlling fungal systemic dissemination (*45*). Neutrophils have shown to be the major immune cells in the control of candidiasis (*67*) by phagocytizing and killing yeast cells and short hyphae, while longer hyphae are killed by inducing neutrophil extracellular traps (NETs), which release DNA, granule enzymes, and antimicrobial peptides (*68*). Our future studies will be focused on understanding the immune cell strategies against fungal CAUTIs and their role in containing the fungal infection in the bladder.

*C. albicans* occupies many niches in the human body, and morphological changes are associated with the establishment of diseased states. This is important in the bladder, since it is an open and dynamic system, where urine is constantly passing through. Therefore, in order to establish a successful colonization, adhesion and biofilm formation on the urinary catheter is essential (*19, 37*). Our results are consistent with that; *Candida* biofilms not only ensure colonization but protect the growing cells from the hostile environment and potentiate the establishment of the infection (*69*). This is the first study unveiling the *C. albicans* mechanism of pathogenesis during CAUTI, finding that Efg1, and specifically its downstream target Als1, are critical for establishing infection (**Fig. 7**). This study opens new avenues to understanding *C. albicans* cellular programming in the catheterized bladder. Importantly, the key interaction of Als1-Fg, an initial step for colonization, could be exploited as a potential therapeutic avenue to prevent *C. albicans* CAUTIs.

**Figure 7.**
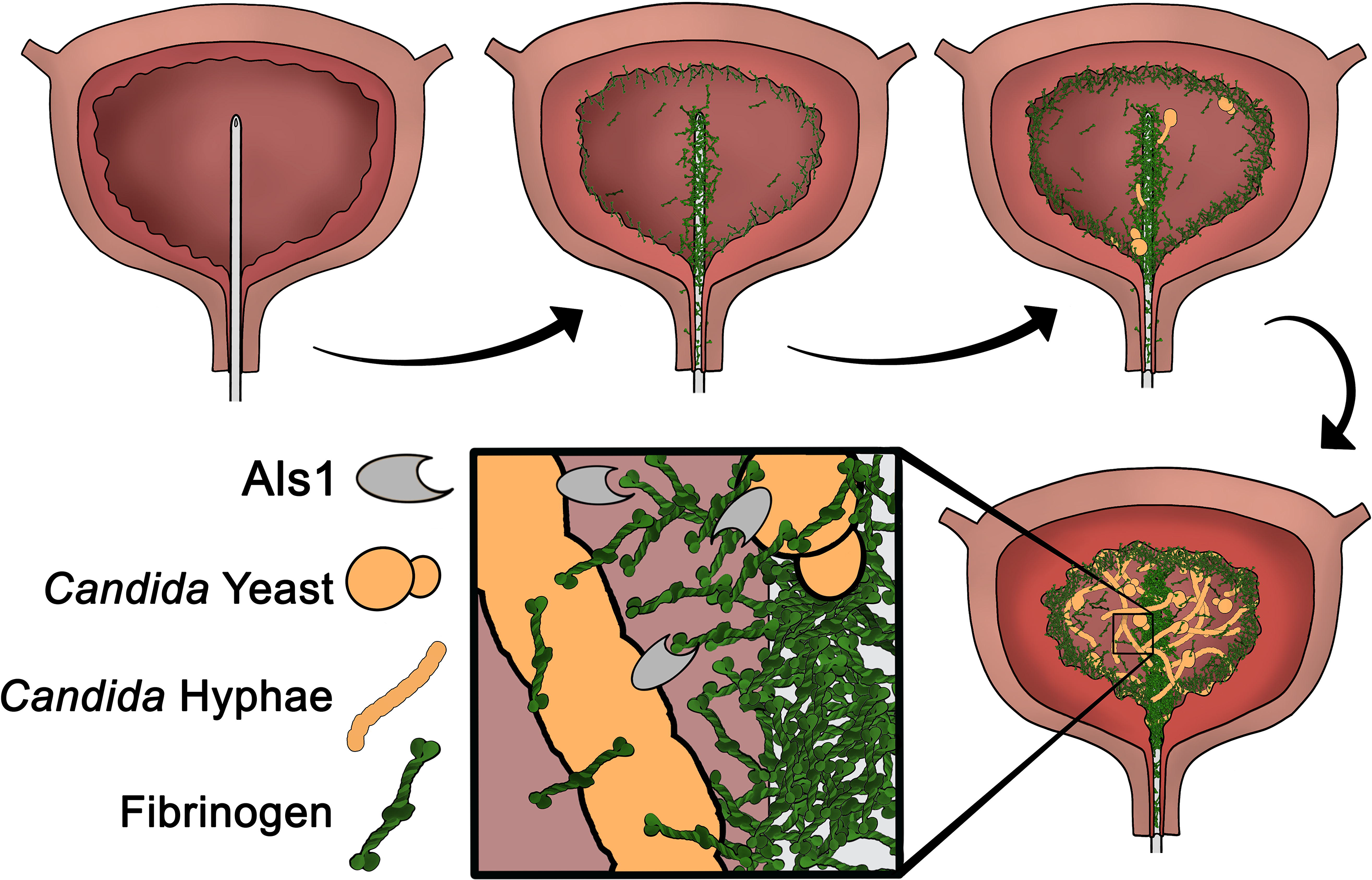
*C. albicans* hyphal morphogenesis and Als1-Fg interactions are critical for CAUTI. Urinary catheterization results in mechanical damage to the bladder and leads to the release of Fg, accumulating in the bladder and depositing on the urinary catheter. The Fg-coated catheter serves as a scaffold for *C. albicans* attachment, leading to robust biofilm formation consisting of yeast, pseudohyphae, and hyphae. *C. albicans* attachment to Fg can be attributed primarily to the adhesin Als1. As a result of this adherence to Fg, *C. albicans* is able to persist in the bladder and establish CAUTI.

## MATERIALS AND METHODS

### Ethics statement

All animal care was consistent with the Guide for the Care and Use of Laboratory Animals from the National Research Council. The University of Notre Dame Institutional Animal Care and Use Committee approved all mouse infections and procedures as part of protocol number 18-08-4792MD. For urine and blood collections, all donors signed an informed consent form and protocols were approved by the Institutional Review Board of the University of Notre Dame under study #19-04-5273 for urine and #18-08-4834 for blood.

### Urine Collection

Human urine from at least two healthy female donors between the ages of 20 - 35 were collected and pooled. Donors did not have a history of kidney disease, diabetes, or recent antibiotic treatment. Urine was sterilized with a 0.22 μm filter (VWR 29186-212) and pH was normalized to 6.0-6.5. BSA (VWR 97061-48); supplemented urine was sterilized again using a 0.22 μm filter. When urine was supplemented with Fg (Enzyme Research Laboratories FIB 3), it was added directly to the sterilized urine and the urine was not sterilized after the addition of Fg.

### Fungal Culture Conditions

All strains of *C. albicans* were cultured at 37 °C with aeration in 5 mL of YPD (10g/L Yeast Extract (VWR J850-500G), 20g/L Peptone (VWR J636-500G), 20 g/L Dextrose (VWR BDH9230-500G)) broth. For *in vivo* mouse experiments, *C. albicans* strains were grown static overnight in 10 mL of YPD.

### Growth Curves

Growth curves were performed in glass test tubes (Thermo Fisher Scientific 14-961-29). Overnight cultures (all in stationary phase; measured using a UV/Vis Spectrophotometer) were normalized to ~1×10^6^ CFU/ml in 1xPBS (Sigma–Aldrich 1002786391). The culture was then diluted (1:1000) into human urine (supplemented with 1 mg/mL BSA, 1 mg/mL Fg, 50X amino acids, or 1 mg/mL human serum), BHI (incubated statically or shaking), or YPD (incubated statically or shaking) and were incubated in the test tube at 37°C for 48 hours. At 0, 24, and 48 hours, samples of each condition were taken and analyzed by CFU counts.

### Morphology Assay and Analysis

All strains of *C. albicans* were grown in YPD with or without serum and in human urine with or without serum. At 0, 24, and 48 hours, a sample of each condition was taken, fixed with 10% formalin, and stained with 100 μg/mL of calcofluor. Samples were viewed under a Zeiss inverted light microscope (Carl Zeiss, Thornwood, NY) with the DAPI fluorescent channel. Random images were taken at 100x magnification and processed with manual counting of yeast, pseudohyphae, and hyphae. Images (consisting of a 3 × 3 tiled region, i.e. 9 fields of view) were randomly acquired and at least three images were analyzed per condition. The total number of cells per phenotype were summed and divided by the total number of cells to give the overall percentage of each cell type.

#### Antibodies and dyes used in this study

##### Primary antibodies

Goat anti-fibrinogen (Sigma-Aldrich F8512), rabbit anti-*Candida* (ThermoFisher Scientific PA1-27158), and rat anti-mouse Ly6G (BD Pharmingen 551459).

##### Secondary antibodies

Alexaflour 488-labeled donkey anti-goat (ThermoFisher Scientific SA5-10086); Alexaflour 594-labeled donkey anti-rabbit (ThermoFisher Scientific SA5-10039); Alexaflour 647-labeled donkey anti-rat (ThermoFisher Scientific SA5-10029); IRDye 800CW donkey anti-goat; and IRDye 680LT donkey anti-rabbit. Alexaflour secondary antibodies were purchased from Invitrogen Molecular Probes and IRDye conjugates secondary antibodies from LI-COR Biosciences.

##### Dyes

Calcofluor white (Fluorescent Brightener 28, Sigma F3543) staining; Hoechst dye (Thermo Fisher Scientific 62249) staining; Hematoxylin and Eosin (H&E) (vector Laboratories #H-3502).

### Biofilm formation assays

Biofilm formation assays were performed in 96 well flat-bottomed plates (VWR 10861-562) coated with 100uL of BSA or Fg (150 μg/mL) incubated overnight at 4°C. The various strains were grown as described above and the inoculum normalized to ~1×10^6^ CFUs/ml. Cultures were then diluted (1:1000) into YPD, BHI, or human urine. 100 μL of the inoculum were incubated in the wells of the 96 well plate at 37°C for 48 hours while static.

Following the 48 hr incubation, the supernatant was removed from the plate and washed three times with 200 μL 1x PBS to remove unbound fungi. Plates were fixed with 10% neutralizing formalin (Leica 3800600) for 20 minutes and followed by three washes with PBS containing 0.05% Tween-20 (PBS-T). Blocking solution (PBS with 1.5% BSA and 0.1% sodium azide (Acros Organics 447811000)) was added to the plate for one hour at room temperature and then washed with PBS-T (3x). Biofilms were incubated with anti-*Candida* antibodies diluted into dilution buffer (PBS with 0.05% Tween (VWR M147-1L), 0.1% BSA) for two hours. Plates were washed three times with PBS-T and incubated for one hour with IRDye 680 LT donkey anti-rabbit secondary antibody solution at room temperature and washed with PBS-T (3x). As a final step, the biofilms were visualized by scanning the plates using the Odyssey Imaging System (LI-COR Biosciences) and the analyzed with Image Studio software to obtain the fluorescence intensities (LI-COR Version 5.2, Lincoln, NE).

### BSA or Fg-coated dishes and formation of Fibrin fibers/nets

For these assays No. 0 cover glass glass-bottom 35 mm petri dish with a 14 mm microwell (MatTek P35G-0-14-C) were used. The dishes were coated with 150 μg/mL of BSA or Fg overnight at 4°C. For fibrin fiber/nets formation, Fg and thrombin (Sigma-Aldrich T6884-250UN) were thawed at 37°C. 100 μl of 0.5 mg/ml Fg in PBS was added into the microwell glass-bottom and then 10 μl of 2 U/ml thrombin was added to polymerize Fg into fibrin. Dishes were incubated at 37°C for 1 hour and kept overnight at 4°C

### Visualization of biofilms and fungal-fibrin interactions

The various strains were grown as described above and the inoculum normalized to ~1×10^6^ CFUs/ml in PBS. These cultures were then diluted (1:1000) into human urine, added to the BSA-, Fg-, or fibrin coated dishes and then were incubated at 37°C for 48 hours under static conditions. After incubation, dishes were then washed three times with 1x PBS to remove unbound fungi, then dishes were fixed with 10% neutralizing formalin solution for 20 minutes and washed with 1x PBS three times. Dishes were blocked with blocking solution for an hour at room temperature as described above. Then BSA- and Fg-coated dishes were incubated in primary antibody (rabbit anti-*Candida*) and for fibrin-coated dishes were incubated with rabbit anti-*Candida* and goat anti-Fg antibodies. Incubation with the primary antibodies was done for two hours followed by three washes with PBS-T. Then, dishes were incubated for 1 hour with Alexaflour 594-labeled donkey anti-rabbit secondary antibody for BSA- and Fg-coated dishes and Alexaflour 594-labeled donkey anti-rabbit and Alexaflour 488-labeled donkey anti-goat antibodies for fibrin-coated dishes, followed by three washes with PBS-T. BSA- and Fg- coated dishes were visualized with a Zeiss inverted light microscope, and images were taken at different magnifications (10x, 20x, 40x and 100x). Zen Pro and Fiji-ImageJ (*70*) softwares were used to analyze the images. For the fungal-fibrin interaction, fibrin-coated dishes were visualized by Nikon A1-R/Multi-Photon Laser Scanning Confocal Microscope and images were analyzed by IMARIS Image Analysis software and ImageJ software (*70*).

### Crystal violet staining

Following biofilm formation on Fg-coated microplate, the supernatant was removed, and the plate was incubated in 200 μl of 0.5% crystal violet for 15 minutes. Crystal violet stain was removed, and the plate was washed with water to remove remaining stain. Plates were dried and then incubated with 200 μl of 33% acetic acid for 15 minutes. In a new plate, 100 μl of the acetic acid solution was transferred, and absorbance values were measured via a plate spectrophotometer at 595 nm.

### *In vivo m*ouse model

Mice used in this study were ~6-week-old female wild-type C57BL/6 mice purchased from Jackson Laboratory. Mice were subjected to transurethral implantation of a silicone catheter and inoculated as previously described (*42*). Briefly, mice were anesthetized by inhalation of isoflurane and implanted with a 6-mm-long silicone catheter (BrainTree Scientific SIL 025). Mice were infected immediately following catheter implantation with 50 μl of ~1◻×◻10^6^ CFUs/mL in PBS, of one of the fungal strains introduced into the bladder lumen by transurethral inoculation. Mice were sacrificed at 24 hours post infection by cervical dislocation after anesthesia inhalation and catheter, bladder, kidneys, spleen, and heart were aseptically harvested. Organs were homogenized and catheters were cut in small pieces before sonication for fungal CFU enumeration. A subset of catheters were fixed for imaging as described below and a subset of bladders were fixed and processed for immunofluorescence and histology analysis as described below.

### Catheter imaging and analysis

Harvested catheters were fixed for imaging via standard IF procedure as previously described (*41*). Briefly, catheters were fixed with formalin, blocked, washed with 1x PBS, and incubated with the appropriate primary antibodies overnight. Catheters were then incubated with secondary antibodies for 2 hrs at room temperature. Catheters were washed with PBS-T and then a final wash with PBS. Catheters were visualized with the Odyssey Imaging System and then analyzed using color pixel counter from Fiji-ImageJ software (*70*). The number of pixels of each color was compared to the total number of pixels to identify percent coverage of the catheter.

### IHC and H&E staining of mouse bladders

Mouse bladders were fixed in 10% formalin overnight, before being processed for sectioning and staining as previously described (*37*). Briefly, bladder sections were deparaffinized, rehydrated, and rinsed with water. Antigen retrieval was accomplished by boiling the samples in Na-citrate, washing in tap water, and then incubating in 1x PBS three times. Sections were then blocked (1% BSA, 0.3% TritonX100 (Acros Organics 21568-2500) in 1x PBS) washed in 1x PBS, and incubated with appropriate primary antibodies diluted in blocking buffer overnight at 4 °C. Next, sections were washed with 1x PBS, incubated with secondary antibodies for 2 hrs at room temperature, and washed once more in 1x PBS prior to Hoechst dye staining. Secondary antibodies for immunohistochemistry were Alexa 488 donkey anti-goat, Alexa 550 donkey anti-rabbit, and Alexa 650 donkey anti-rat. Hematoxylin and Eosin (H&E) stain for light microscopy was done by the CORE facilities at the University of Notre Dame (ND CORE). All imaging was done using a Zeiss inverted light microscope and a Nikon A1-R/Multi-Photon Laser Scanning Confocal Microscope. Zen Pro, ImageJ software, and IMARIS Image Analysis software were used to analyze the images.

### Measurement of *EFG1* and *ALS1* expression levels during *C. albicans* CAUTI

Total RNA was extracted from infected and catheterized mouse bladders with the Qiagen RNeasy Kit (Cat. No. 74004) and subsequently DNaseI treated. RNA was reverse transcribed into cDNA via qScript cDNA synthesis kit (QuantaBio, Cat. No.101414) and incubated for 5 minutes at 25°C, 30 minutes at 42°C, 5 minutes at 85°C, and held at 4°C. qPCR was performed on QuantStudio 3 (Thermo Fisher Scientific) under the following conditions: 95°C^5min^ (95^15sec^, 60^60sec^)_35 cycles_ (95^15sec^, 60^60sec^, 95^15sec^) _melt curve_. Data were normalized using *ITS2* as an internal housekeeping control gene (*71*) and WT as a sample calibrator and set the expression to 100%. Data were analyzed via the ΔΔCT method. Primers are described in **Supplementary Table 4**.

### Neutrophil analysis

Mouse bladders were harvested and minced into digestion buffer (34U/mL Liberase and 100 μg/mL DNase1 in D-PBS). Bladder were incubated for 1 hours in a heat block shaker at 37°C and 250rpm and every 15 min were vortexed in high speed for 30s. Digestion was arrested by addition of D-PBS supplemented with 2% FBS and 0.2 μM EDTA and passed through 40-μm cell strainers. Following a 20-minute incubation in Fc block (BD Biosciences), cells were immunolabeled with the antibodies listed in **Supplementary Table 5**. Total cell counts in the bladder were determined on a BD LSRFortessa X-20 system cytometer (using DIVA software and analyzed by FlowJo software).

### Statistical analysis

Data from at least 3 experiments were pooled for each assay. Two-tailed Mann-Whitney *U* tests were performed with GraphPad Prism 5 software (GraphPad Software, San Diego, CA) for all comparisons described in biofilm, CAUTI, and catheter coverage experiments. Values represent means ± SEM derived from at least 3 independent experiments. *, *P*<0.05; **, *P*<0.005; ***, *P*<0.0005; ns, difference not significant.

## Acknowledgments

We thank members of the A.L.F.M. and F.H.S.T. laboratories for their helpful suggestions and insightful comments. Special thank you to Dr. Michael G. Caparon for his comments. Special thank you to Dr. Sara Cole and the ND Integrated Imaging Facility for tissue processing and support during imaging.

## Author contributions

A.L.F.M., and F.H.S.T. designed the experiments. A.A.L., M.J.A., A.M., J.J.M., and P.S. performed the studies. R.S.S., N.C.G., and L.W. designed and generated the adhesin mutant strain collection and the *ALS1* OE strain. C.J.N. generated the *efg1*Δ/Δ, *EFG1* complemented, *als1*Δ/Δ, and *ALS1* complemented strains. A.A.L., A.L.F.M, and F.H.S.T. wrote the paper. All authors contributed to editing the paper.

