## Supplemental figures for "The catheterized bladder environment promotes Efg1- and Als1-dependent *Candida albicans* infection"

**Supplementary Figures**.

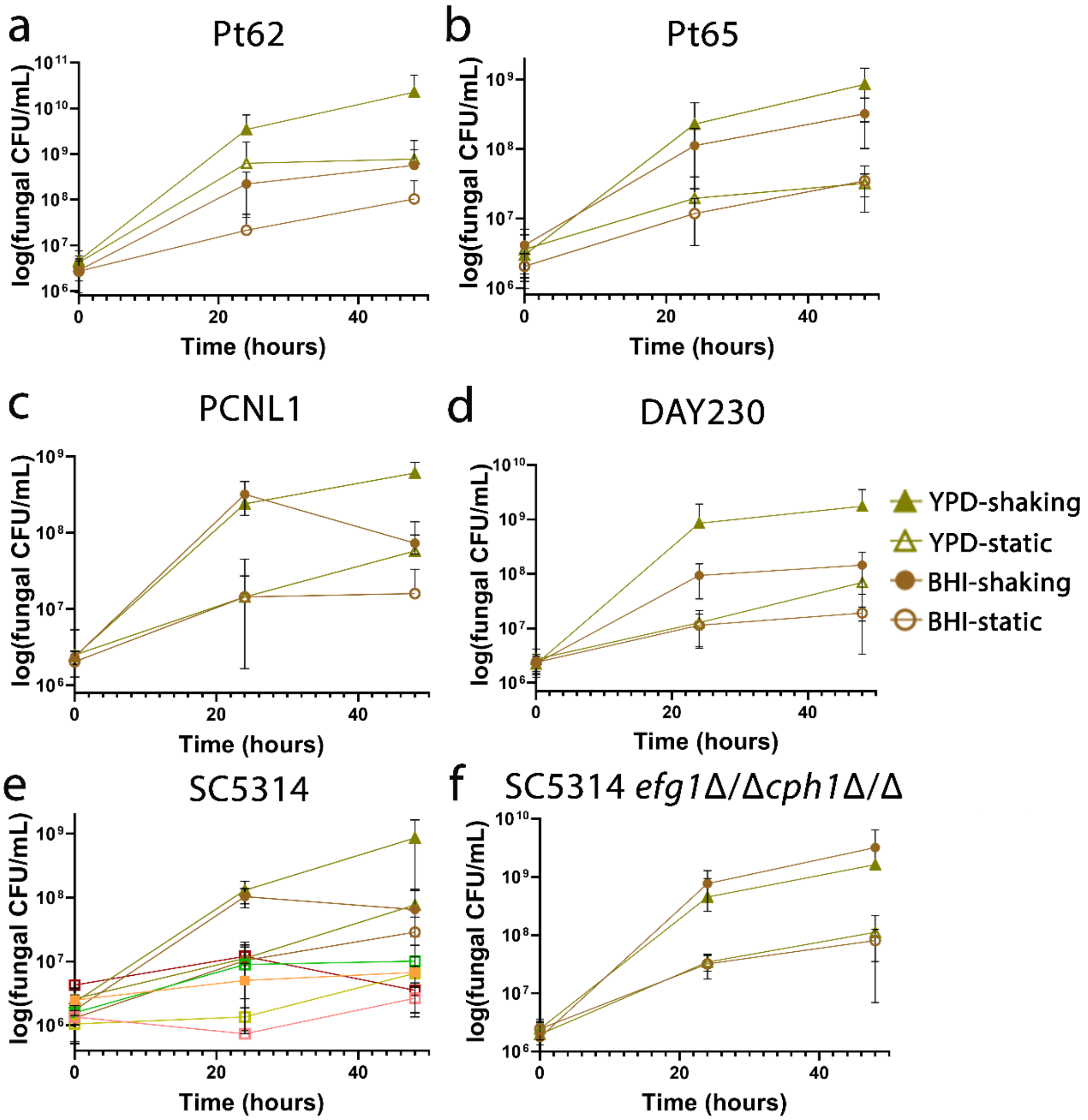


**Supplementary Figure 1. *C. albicans* growth in rich media. (A-F)** Growth curves of *C. albicans* strains Pt62 (A), Pt65 (B), PCNL1 (C), DAY286 (D), SC5314 (E), and SC5314 *efg1*Δ/Δ*cph1*Δ/Δ (F), grown in n YPD, and BHI media. Fungal growth was determined by CFU enumeration after 0, 24, and 48 hours under shaking and static conditions. Data presented shows the mean and standard error of the mean derived from three independent experiments with at least three technical replicates.

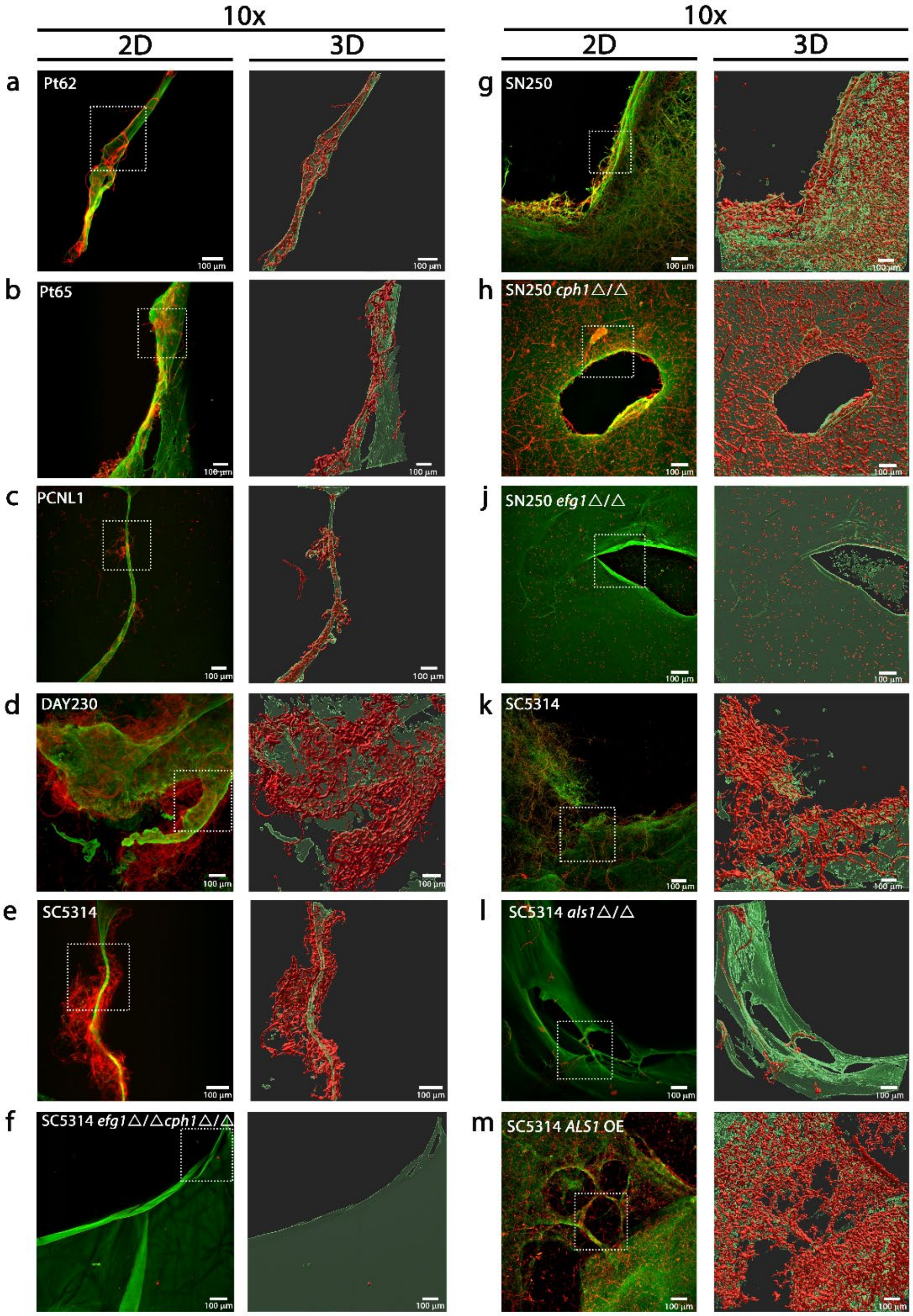


**Supplementary Figure 2.** ***C. albicans*-interaction with fibrin fibers/nets in urine conditions.** (A-F) Microscopically visualization and 3D reconstruction of 48 hrs *C. albicans* biofilms on fibrin fibers/nets grown in human urine using antibodies against Fg (anti-Fg; green) and *C. albicans* (anti-*Candida*; red). Scale bars: 100 µm for 10x. White squares represent zoomed-in areas of higher magnification (40x) shown in **Fig. 2, 5, and 6**.

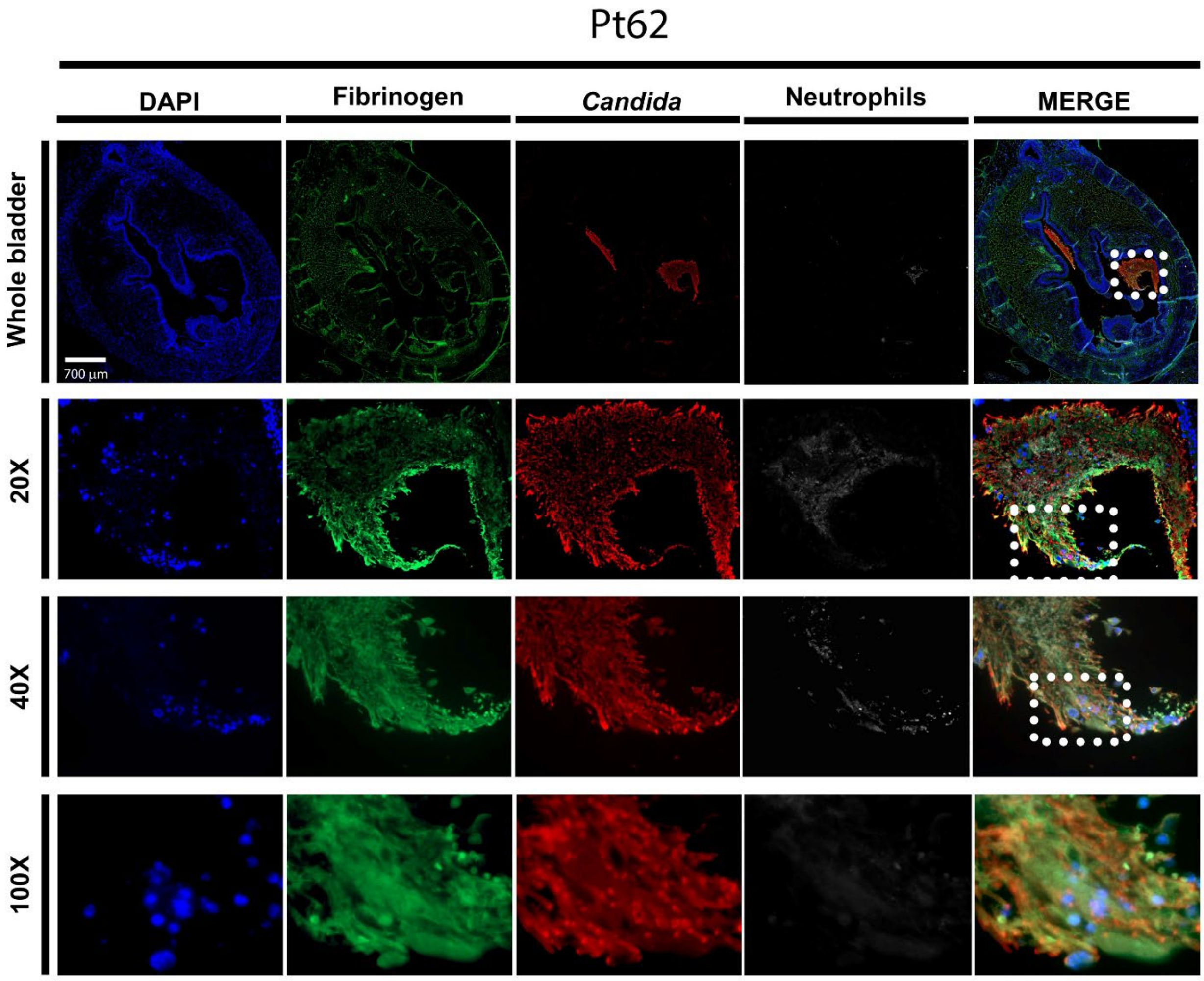


**Supplementary Figure 3.** ***C. albicans*** **Pt62 bladder colonization during CAUTI.** Mice were implanted and infected with 1x $10^6$ CFUs of strain Pt62. At 24 hpi, bladder tissues were harvested, fixed, and parafilm-embedded. Bladders were subjected to IF analysis using antibodies to detect Fg (anti-Fg; green), *C. albicans* (anti-*Candida*; red), and neutrophils (anti-Ly6G; white). Staining with DAPI (blue) delineated the urothelium and cell nuclei (representative images). IF stained bladder scale bars, 700 µm. Magnification (20x, 40x, and 100x).

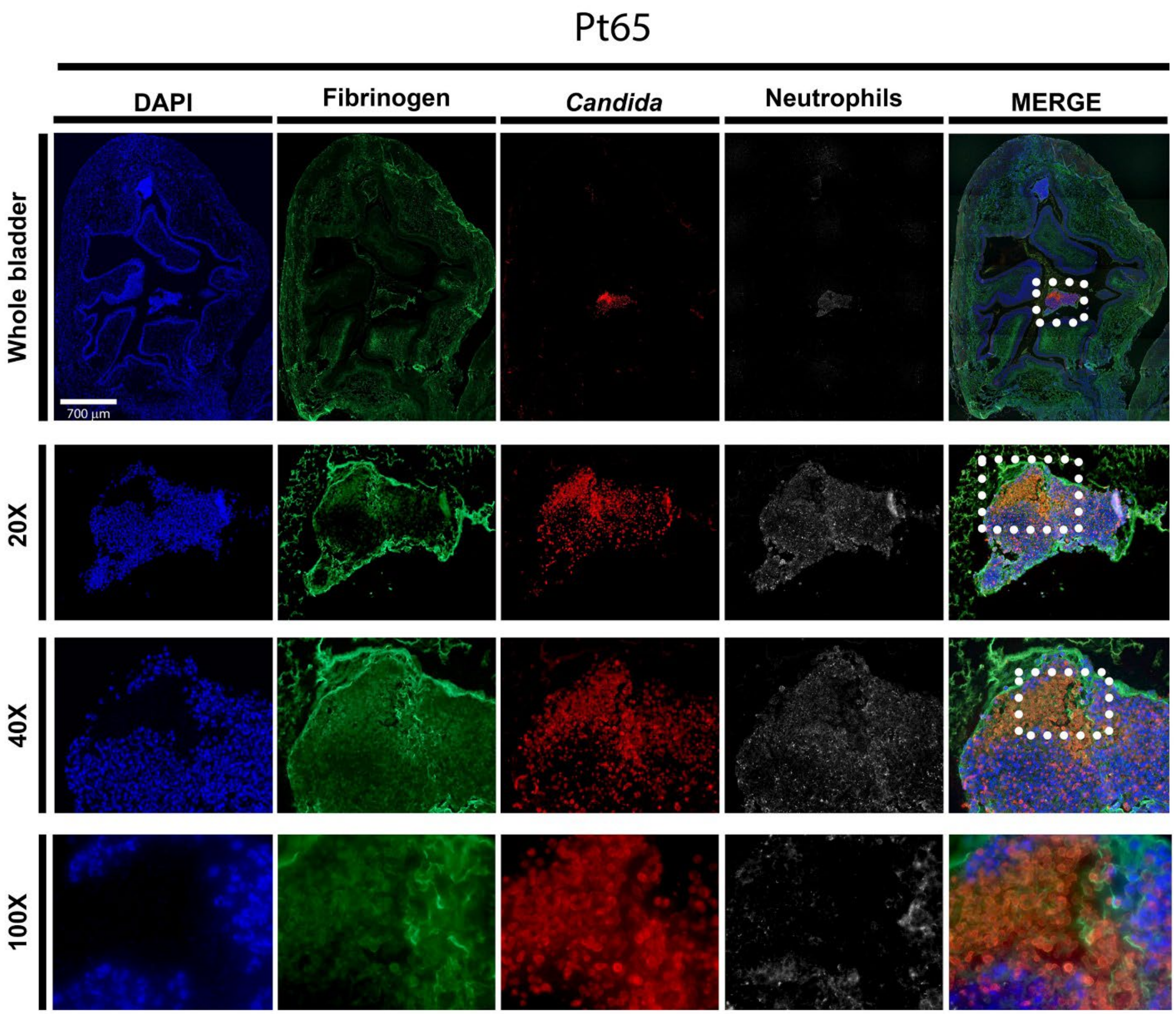


**Supplementary Figure 4.** ***C. albicans*** **Pt65 bladder colonization during CAUTI.** Mice were implanted and infected with 1x $10^6$ CFUs with strain Pt65. At 24 hpi, bladder tissues were harvested, fixed, and parafilm-embedded. Bladders were subjected to IF analysis using antibodies to detect Fg (anti-Fg; green), *C. albicans* (anti-*Candida*; red), and neutrophils (anti-Ly6G; white). Staining with DAPI (blue) delineated the urothelium and cell nuclei (representative images). IF stained bladder scale bars, 700 µm. Magnification (20x, 40x, and 100x).

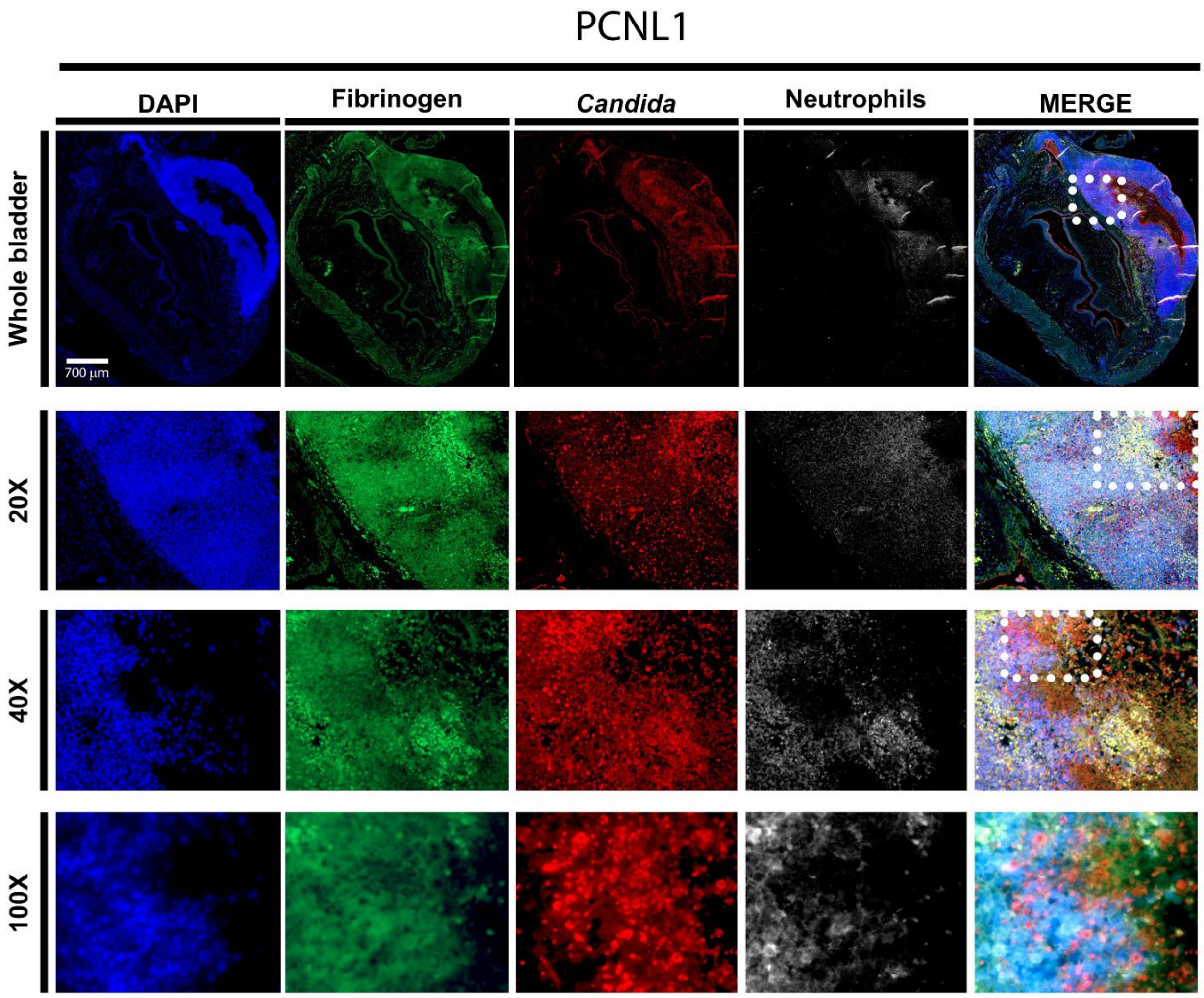


**Supplementary Figure 5. *C. albicans* PCNL1 bladder colonization during CAUTI.** Mice were implanted and infected with 1x $10^6$ CFUs with strain PCNL. At 24 hpi, bladder tissues were harvested, fixed, and parafilm-embedded. Bladders were subjected to IF analysis using antibodies to detect Fg (anti-Fg; green), *C. albicans* (anti-*Candida*; red), and neutrophils (anti-Ly6G; white). Staining with DAPI (blue) delineated the urothelium and cell nuclei (representative images). IF stained bladder scale bars, 700 µm. Magnification (20x, 40x, and 100x).

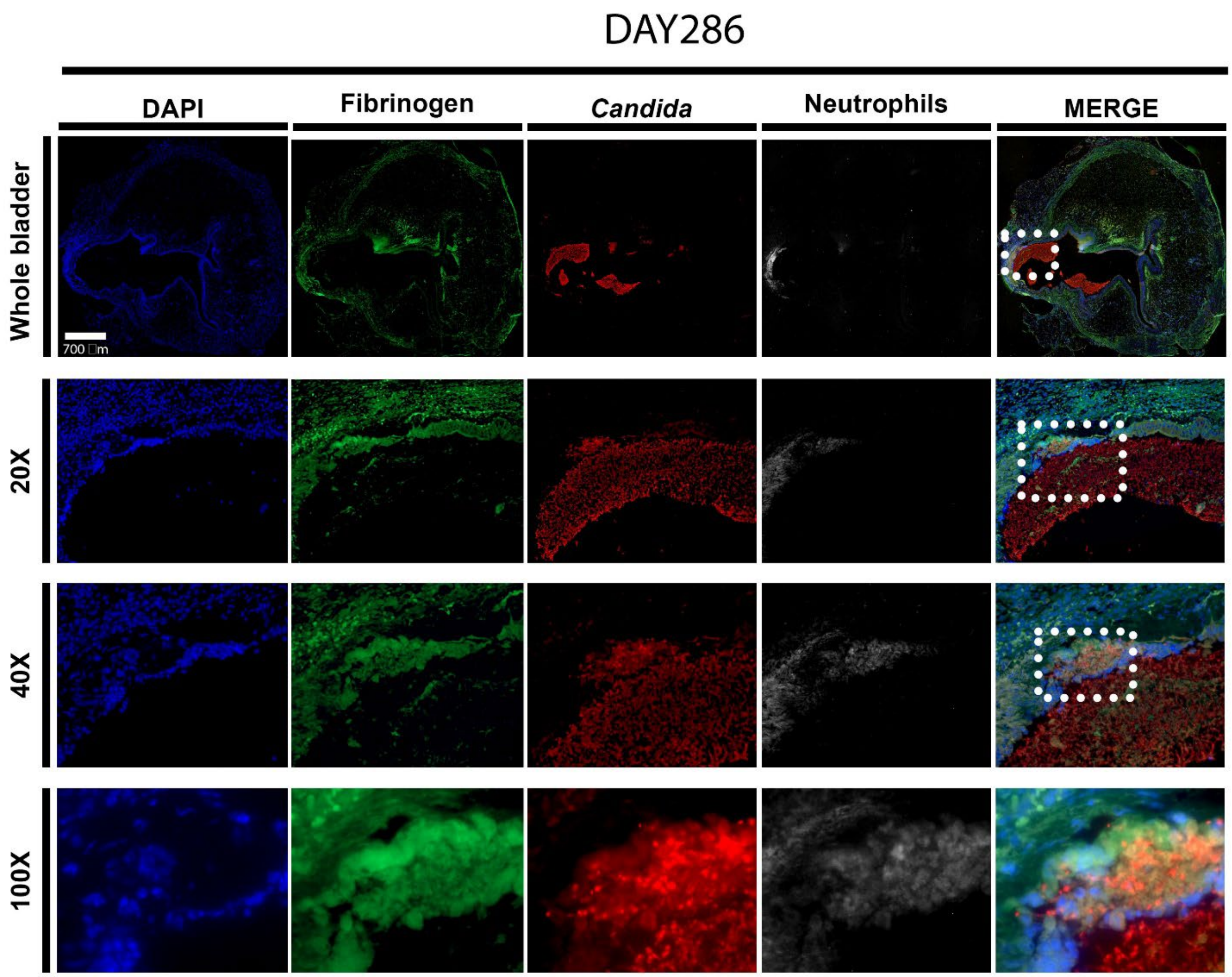


**Supplementary Figure 6. *C. albicans* DAY286 bladder colonization during CAUTI.** Mice were implanted and infected with 1x $10^6$ CFUs with strain DAY286. At 24 hpi, bladder tissues were harvested, fixed, and parafilm-embedded. Bladders were subjected to IF analysis using antibodies to detect Fg (anti-Fg; green), *C. albicans* (anti-*Candida*; red), and neutrophils (anti-Ly6G; white). Staining with DAPI (blue) delineated the urothelium and cell nuclei (representative images). IF stained bladder scale bars, 700 μm. Magnification (20x, 40x, and 100x).

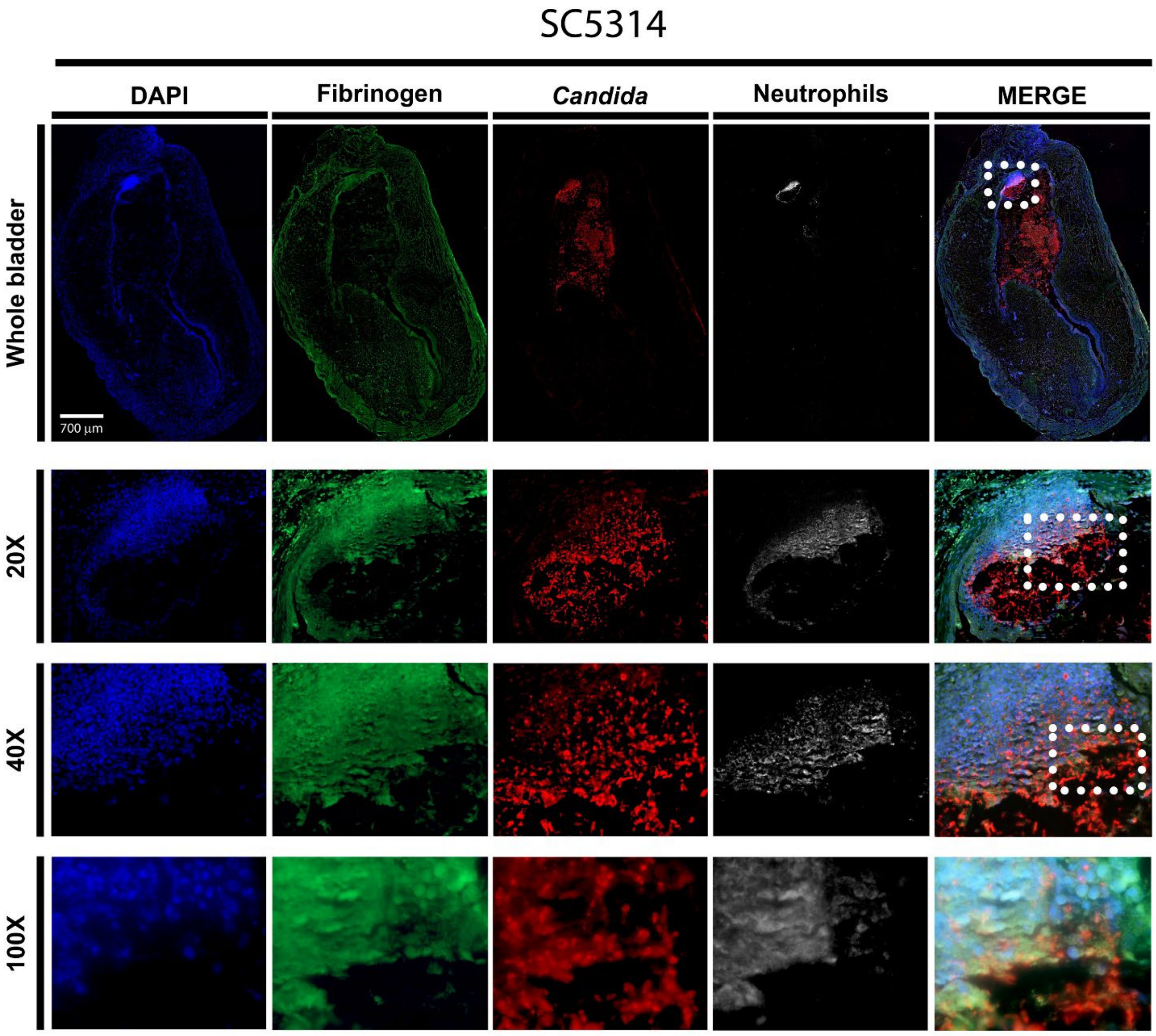


**Supplementary Figure 7. *C. albicans* SC5314 bladder colonization during CAUTI.** Mice were implanted and infected with 1x $10^6$ CFUs with strain SC5314. At 24 hpi, bladder tissues were harvested, fixed, and parafilm-embedded. Bladders were subjected to IF analysis using antibodies to detect Fg (anti-Fg; green), *C. albicans* (anti-*Candida*; red), and neutrophils (anti-Ly6G; white). Staining with DAPI (blue) delineated the urothelium and cell nuclei (representative images). IF stained bladder scale bars, 700 μm. Magnification (20x, 40x, and 100x).

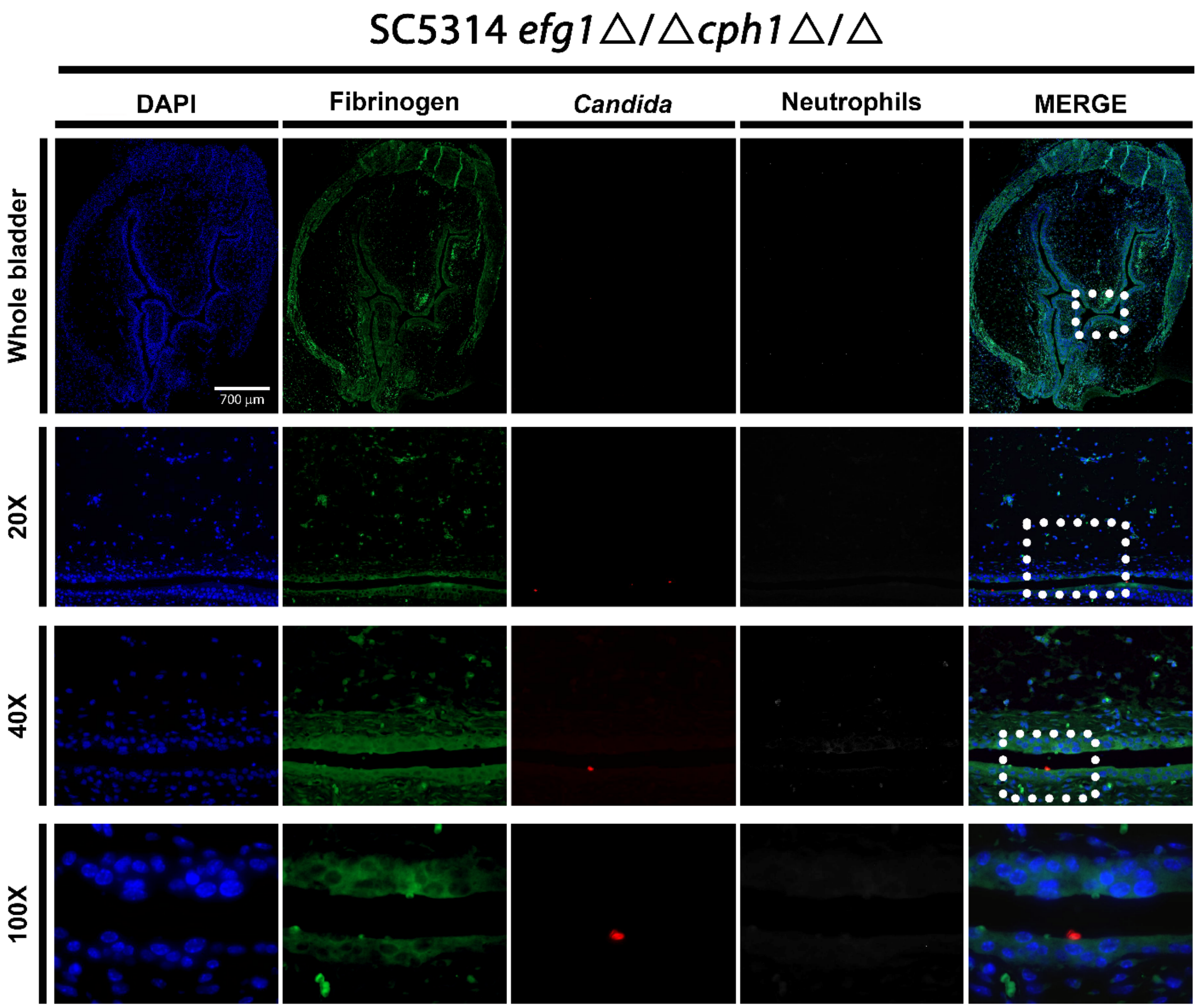


**Supplementary Figure 8.** ***C. albicans*** **SC5314** ***efg1*****Δ/Δ*****cph1*****Δ/Δ bladder colonization during CAUTI.** Mice were implanted and infected with 1x $10^6$ CFUs with strain SC5314 *efg1*Δ/Δ*cph1*Δ/Δ. At 24 hpi, bladder tissues were harvested, fixed, and parafilm-embedded. Bladders were subjected to IF analysis using antibodies to detect Fg (anti-Fg; green), *C. albicans* (anti-*Candida*; red), and neutrophils (anti-Ly6G; white). Staining with DAPI (blue) delineated the urothelium and cell nuclei (representative images). IF stained bladder scale bars, 700 µm. Magnification (20x, 40x, and 100x).

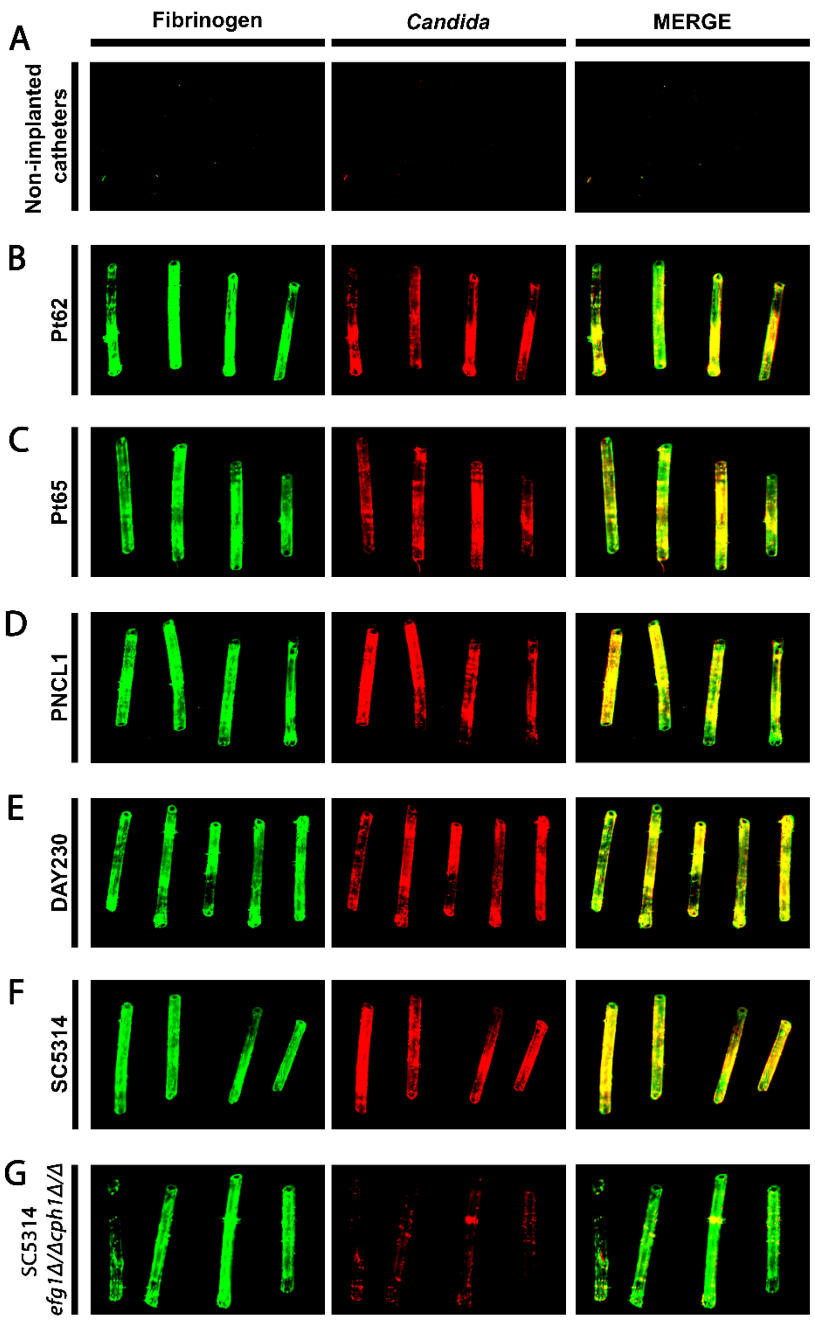


**Supplementary Figure 9. Colocalization of *C. albicans* strains with deposited fibrinogen on catheters during CAUTI.** Catheterized mice were challenged with $1x10^6$ CFUs of the indicated *C. albicans* strain. Implanted catheters were retrieved 24 hpi and stained with antibodies to detect Fg (anti-Fg; green) and *C. albicans* (anti-*Candida*; red) (**B-G**). Non-implanted catheters were used as negative controls (**A**).

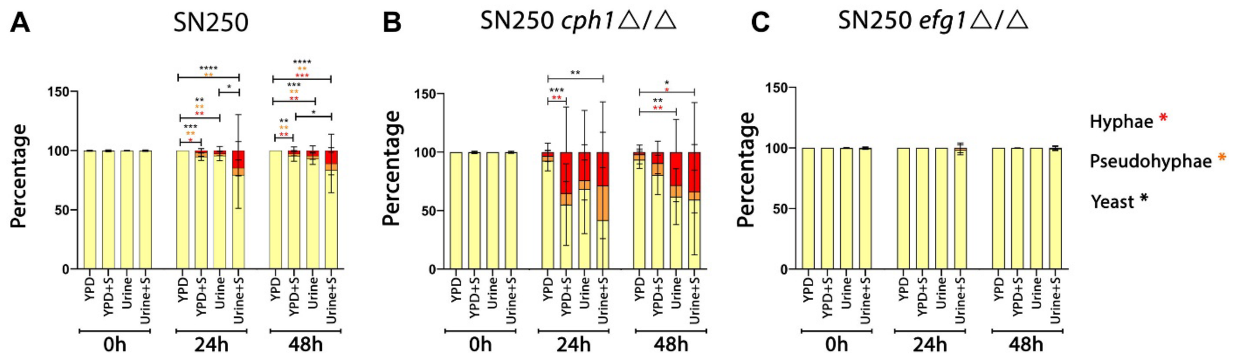


**Supplementary Figure 10. Urine conditions prompt hyphal formation in the *cph1*Δ/Δ strain but not in the *efg1*Δ/Δ strain.** The morphology of *C. albicans* strains were evaluated after 0, 24, and 48 hours of growth in urine and YPD with or without 10% human serum. Representative images were taken at 100x magnification and processed with manual counting of yeast, pseudohyphae, and hyphae. Images (consisting of a 3 x 3 tiled region, i.e. 9 fields of view) were randomly acquired and at least three images were analyzed per condition. The total number of cells per phenotype were summed and divided by the total number of cells to give the overall percentage of each cell type.

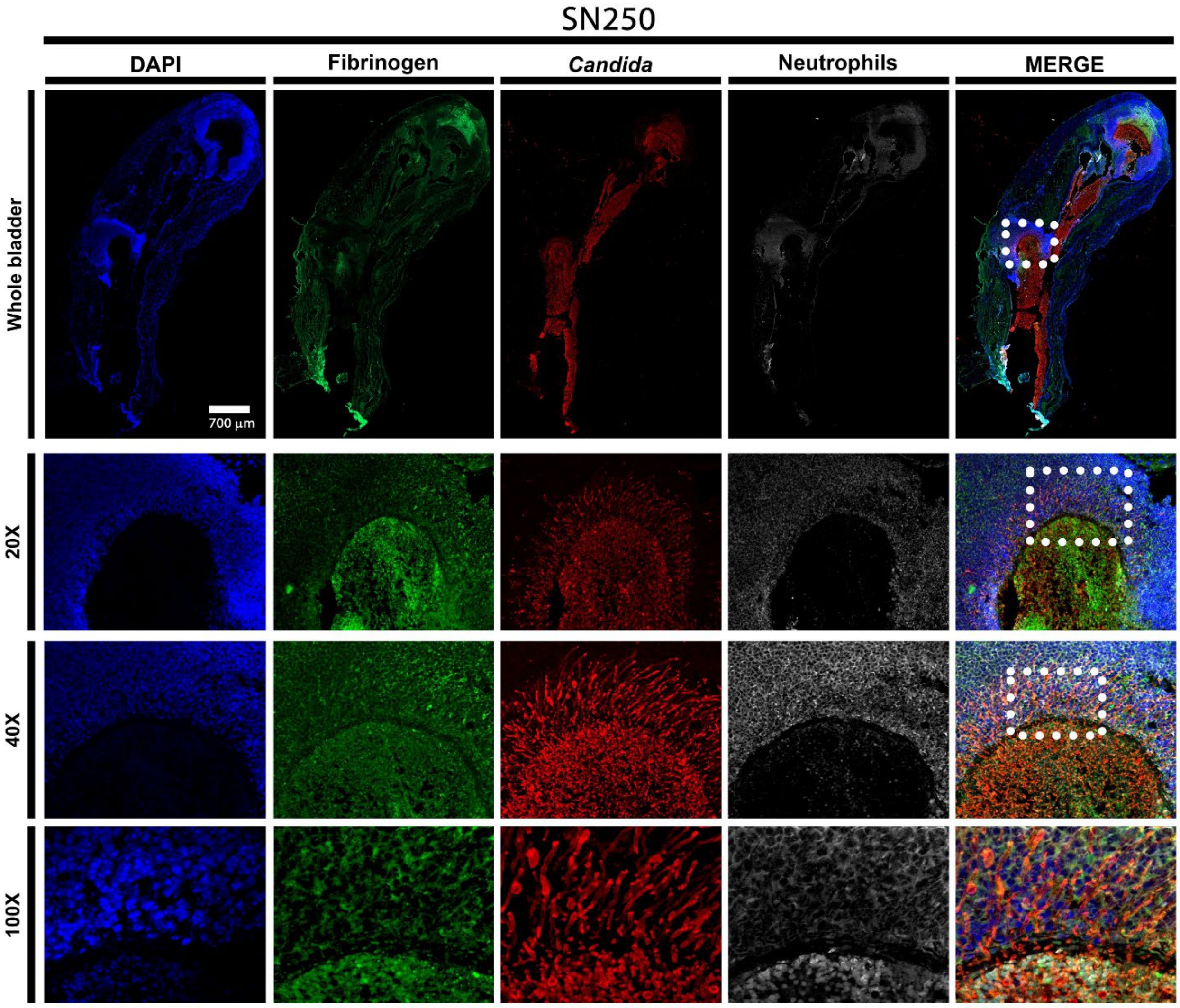


**Supplementary Figure 11. *C. albicans* SN250 bladder colonization during CAUTI.** Mice were implanted and infected with 1x $10^6$ CFUs with strain SN250. At 24 hpi, bladder tissues were harvested, fixed, and parafilm-embedded. Bladders were subjected to IF analysis using antibodies to detect Fg (anti-Fg; green), *C. albicans* (anti-*Candida*; red), and neutrophils (anti-Ly6G; white). Staining with DAPI (blue) delineated the urothelium and cell nuclei (representative images). IF stained bladder scale bars, 700 µm. Magnification (20x, 40x, and 100x). White squares represent zoomed-in areas of higher magnification (20x, 40x, and 100x).

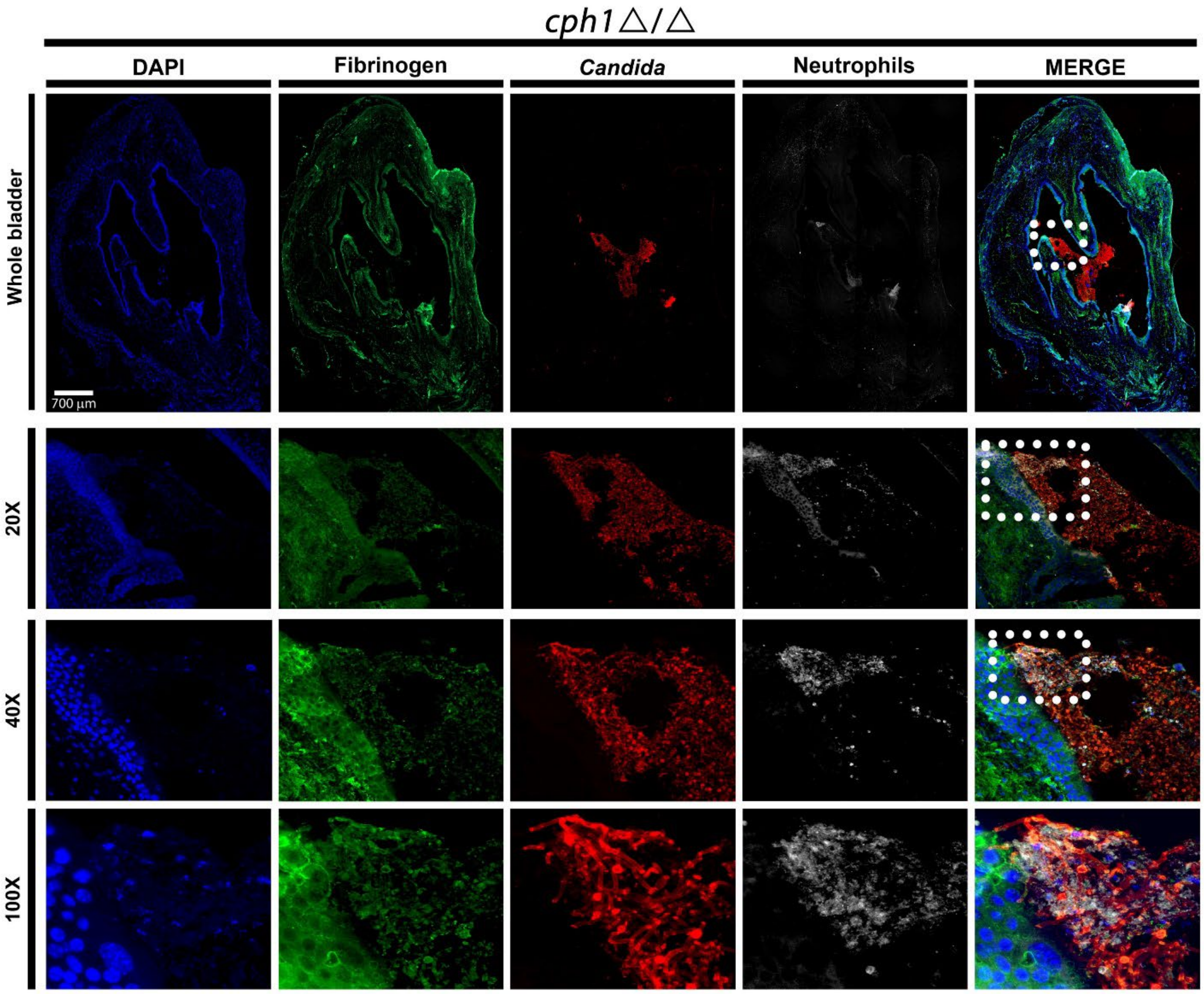


**Supplementary Figure 12.** ***C. albicans*** **SN250** ***cph1*****Δ/Δ bladder colonization during CAUTI.** Mice were implanted and infected with 1x $10^6$ CFUs with strain SN250 *cph1*Δ/Δ. At 24 hpi, bladder tissues were harvested, fixed, and parafilm-embedded. Bladders were subjected to IF analysis using antibodies to detect Fg (anti-Fg; green), *C. albicans* (anti-*Candida*; red), and neutrophils (anti-Ly6G; white). Staining with DAPI (blue) delineated the urothelium and cell nuclei (representative images). IF stained bladder scale bars, 700 µm. White squares represent zoomed-in areas of higher magnification (20x, 40x, and 100x).

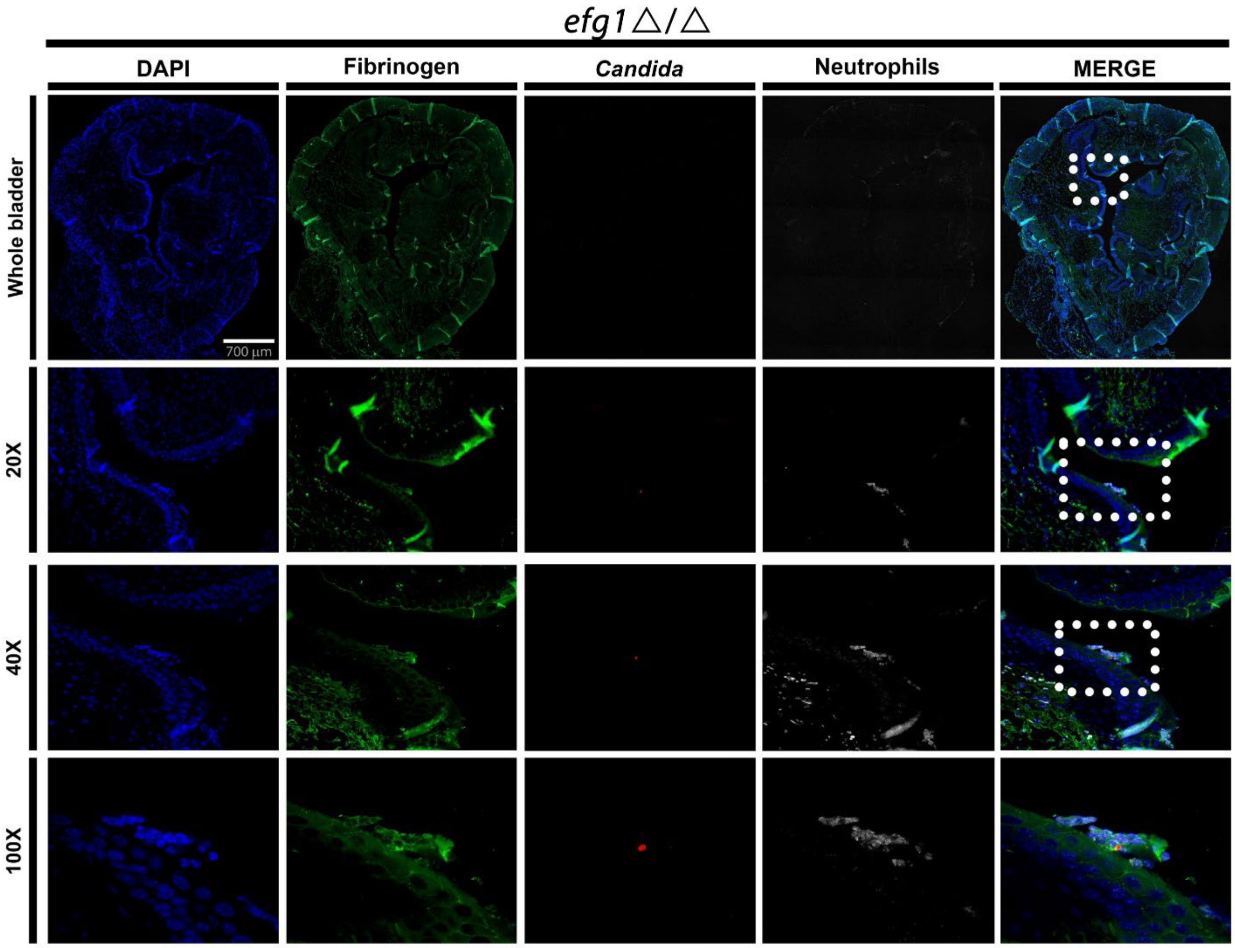


**Supplementary Figure 13.** ***C. albicans*** **SN250** ***efg1*****Δ/Δ bladder colonization during CAUTI.** Mice were implanted and infected with 1x $10^6$ CFUs with strain SN250 *efg1*Δ/Δ. At 24 hpi, bladder tissues were harvested, fixed, and parafilm-embedded. Bladders were subjected to IF analysis using antibodies to detect Fg (anti-Fg; green), *C. albicans* (anti-*Candida*; red), and neutrophils (anti-Ly6G; white). Staining with DAPI (blue) delineated the urothelium and cell nuclei (representative images). IF stained bladder scale bars, 700 µm. White squares represent zoomed-in areas of higher magnification (20x, 40x, and 100x).

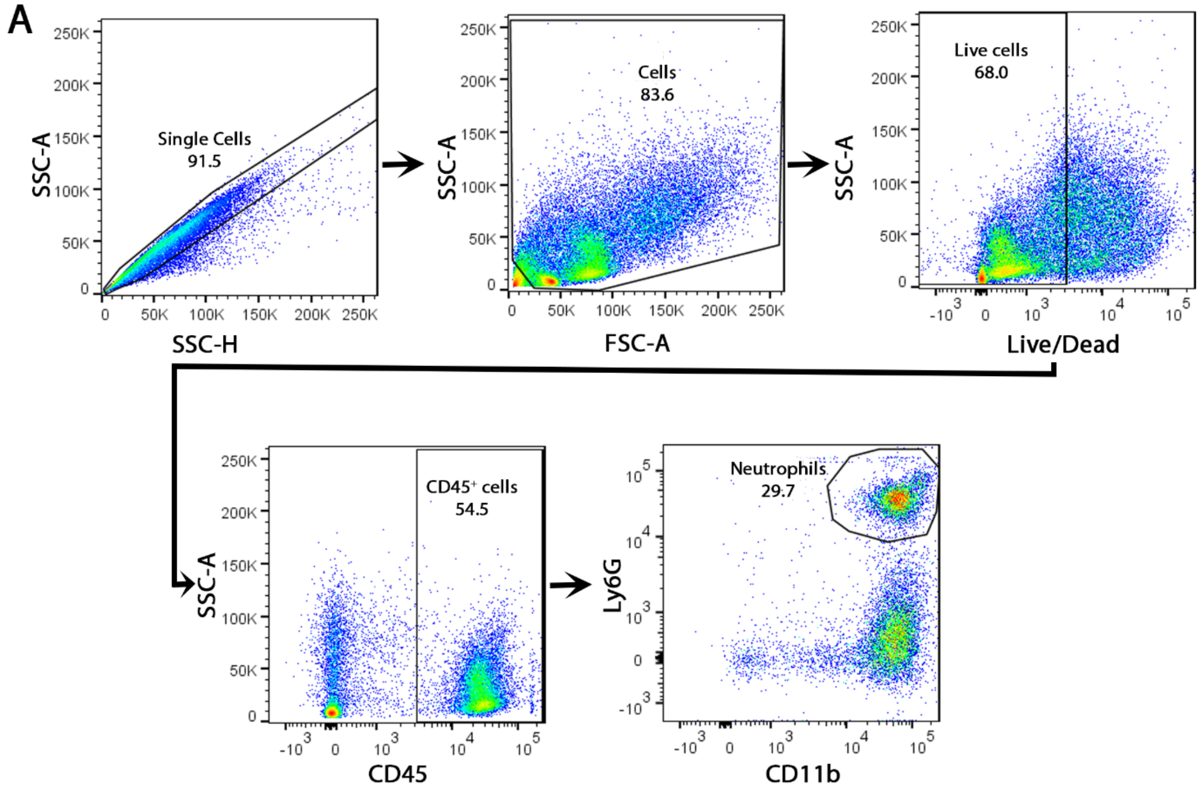


**Supplementary Figure 14. Analysis of neutrophil population in the bladder.** Flow cytometry gating strategy to identify neutrophils.

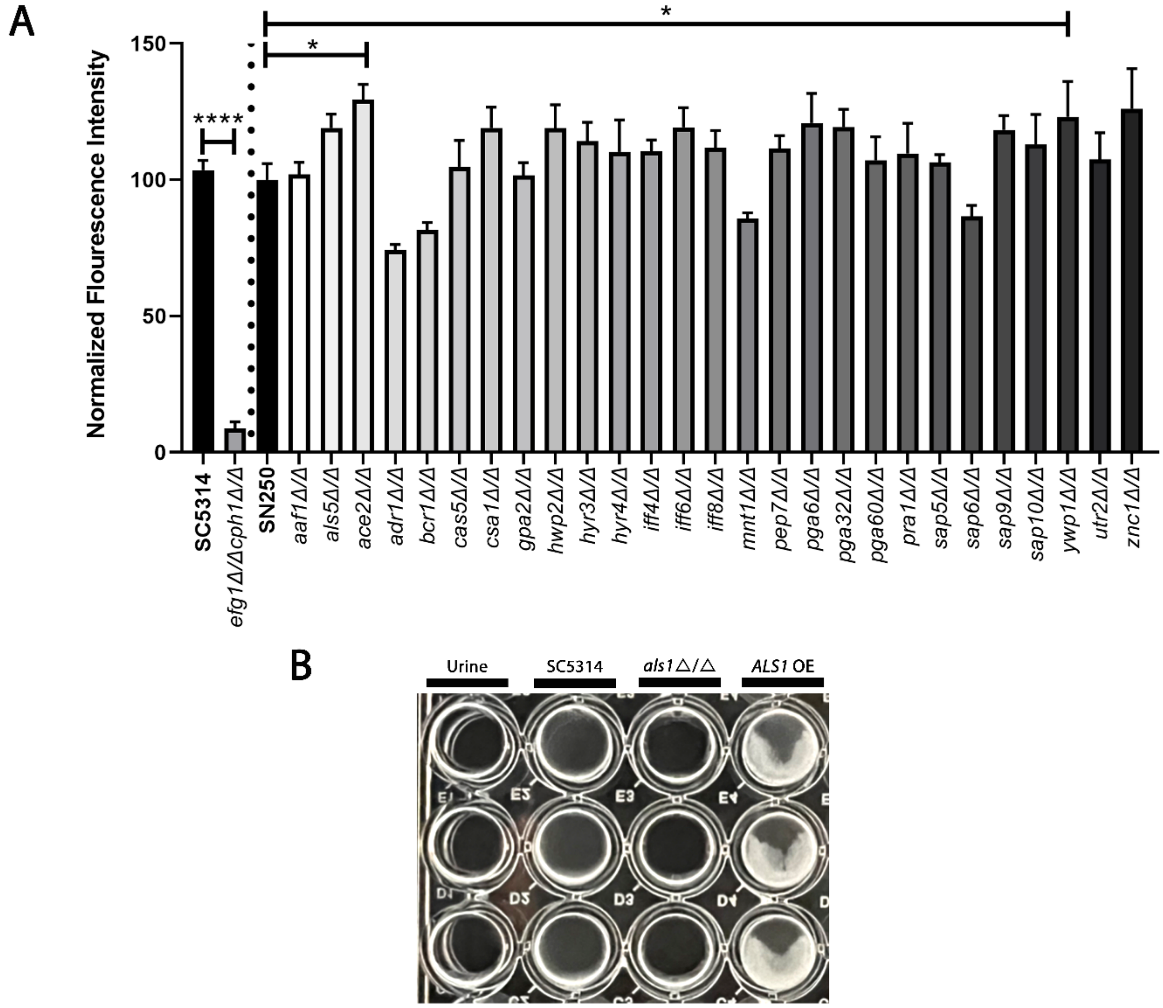


**Supplementary Figure 15. Fungal adhesins contributing to Fg-dependent biofilm formation.** **(A)** Immunostaining analysis of biofilm formation on Fg-coated microplates by *C. albicans* adhesin mutant strains when grown in human urine. At 48 hrs, *C. albicans* biofilm formation was measured by fluorescence intensity using anti-*Candida* antibodies. Data presented shows the mean and standard error derived from three independent experiments with 24 technical replicates. Differences between groups were tested for significance using the Mann-Whitney U test. *, $P < 0.05$. **(B)** Visualization of fungal biofilm biomass on Fg-coated microplates under urine conditions at 48 hrs.

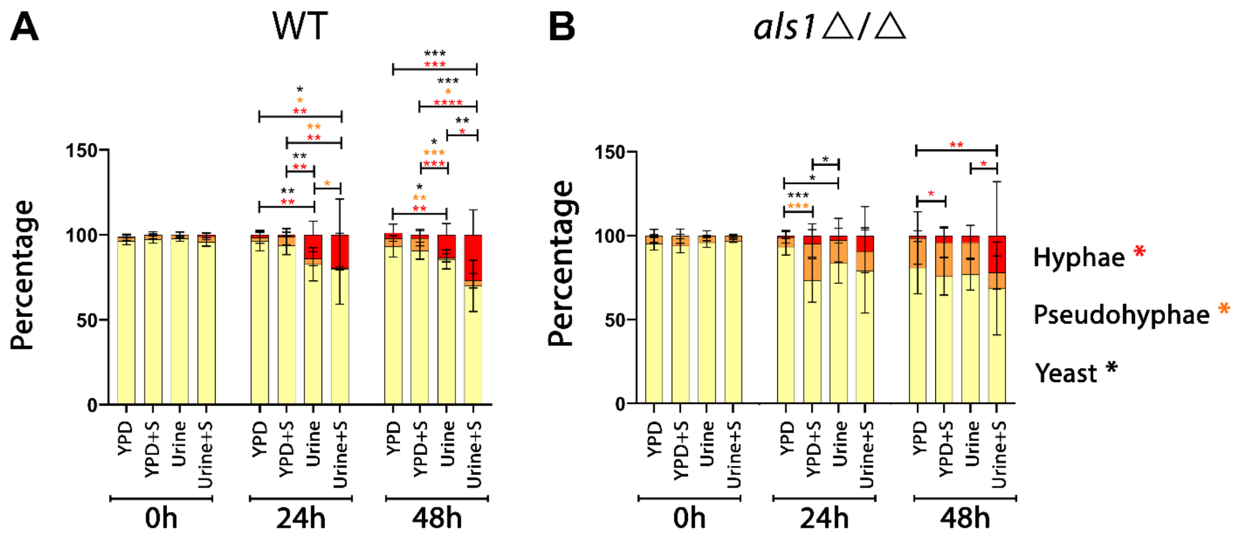


**Supplementary Figure 16. Urine conditions prompt hyphal formation in *als1*Δ/Δ mutant strain similar to the WT strain.** The morphology of *C. albicans* strains were evaluated after 0, 24, and 48 hours of growth in urine and YPD with or without 10% human serum. Random images were taken at 100x magnification and processed with manual counting of yeast, pseudohyphae, and hyphae. Images (consisting of a 3 x 3 tiled region, i.e. 9 fields of view) were randomly acquired and at least three images were analyzed per condition. The total number of cells per phenotype were summed and divided by the total number of cells to give the overall percentage of each cell type.

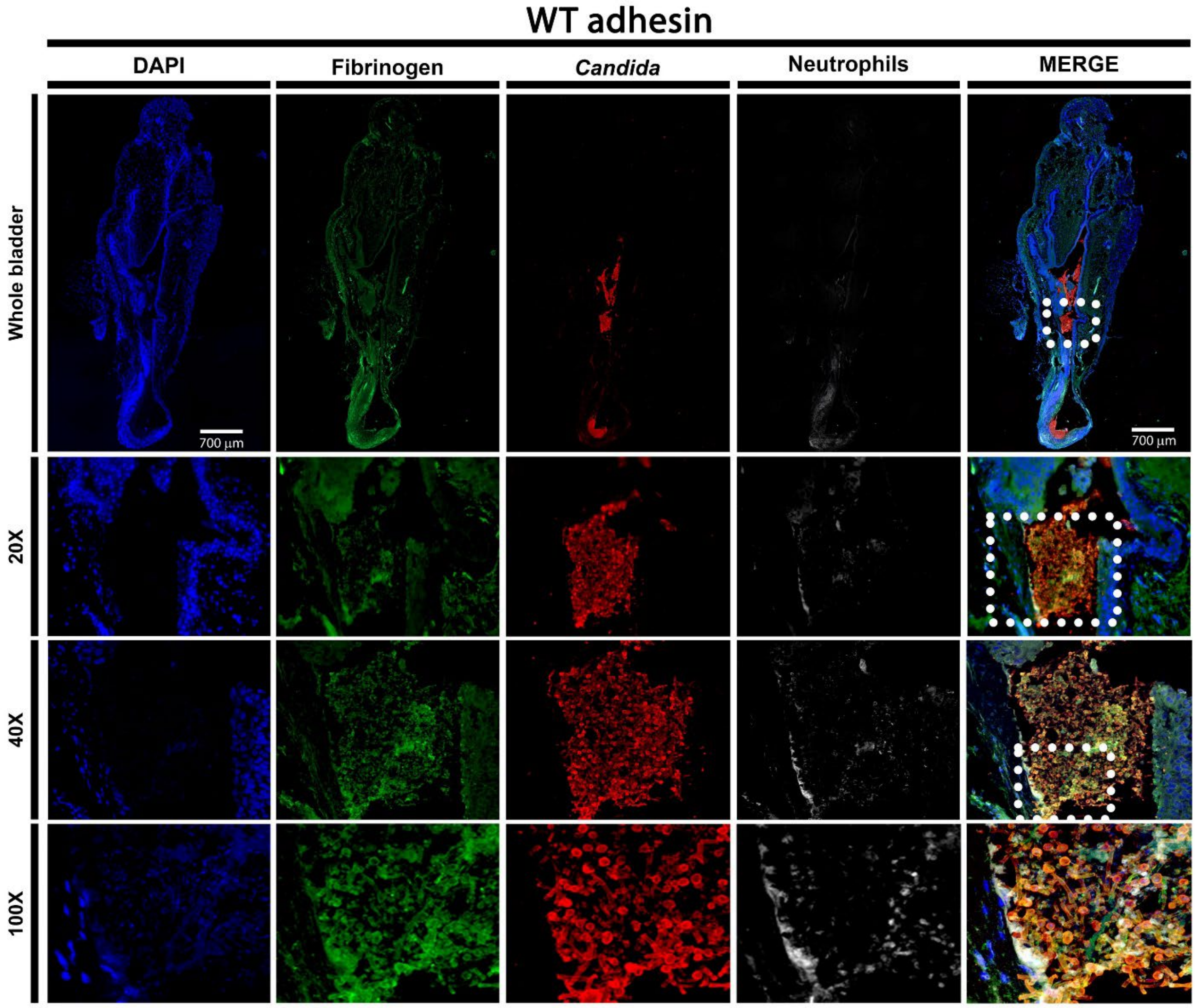


**Supplementary Figure 17.** ***C. albicans*** **WT adhesin bladder colonization during CAUTI.** Mice were implanted and infected with 1x $10^6$ CFUs with the WT adhesin strain. At 24 hpi, bladder tissues were harvested, fixed, and parafilm-embedded. Bladders were subjected to IF analysis using antibodies to detect Fg (anti-Fg; green), *C. albicans* (anti-*Candida*; red), and neutrophils (anti-Ly6G; white). Staining with DAPI (blue) delineated the urothelium and cell nuclei (representative images). IF stained bladder scale bars, 700 μm. White squares represent zoomed-in area of higher magnification (20x, 40x, and 100x).

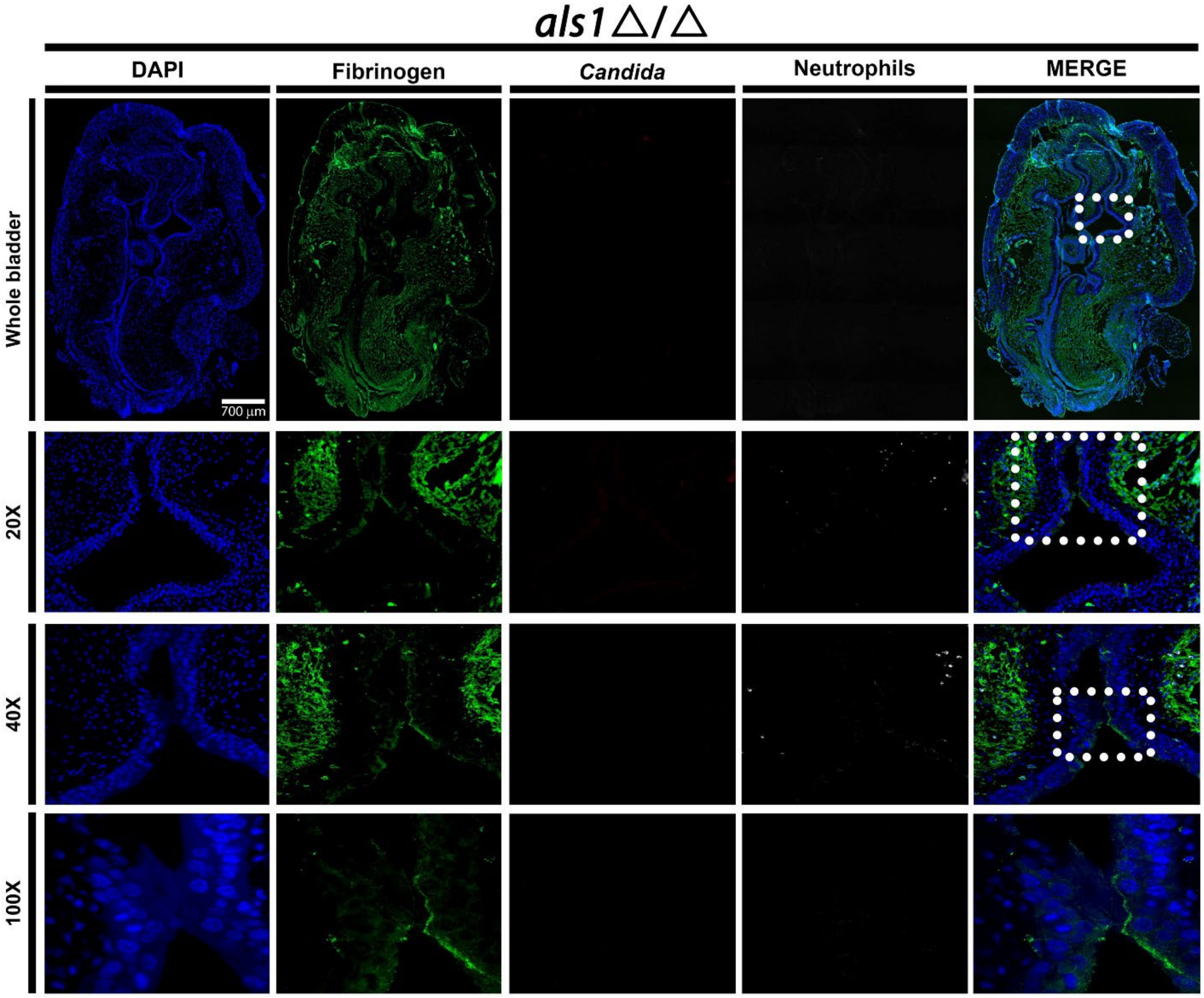


**Supplementary Figure 18.** ***C. albicans*** **SC5314** ***als1*****Δ/Δ bladder colonization during CAUTI.** Mice were implanted and infected with 1x $10^6$ CFUs with strain SC5314 *als1*Δ/Δ. At 24 hpi, bladder tissues were harvested, fixed, and parafilm-embedded. Bladders were subjected to IF analysis using antibodies to detect Fg (anti-Fg; green), *C. albicans* (anti-*Candida*; red), and neutrophils (anti-Ly6G; white). Staining with DAPI (blue) delineated the urothelium and cell nuclei (representative images). IF stained bladder scale bars, 700 μm. White squares represent zoomed-in areas of higher magnification (20x, 40x, and 100x).

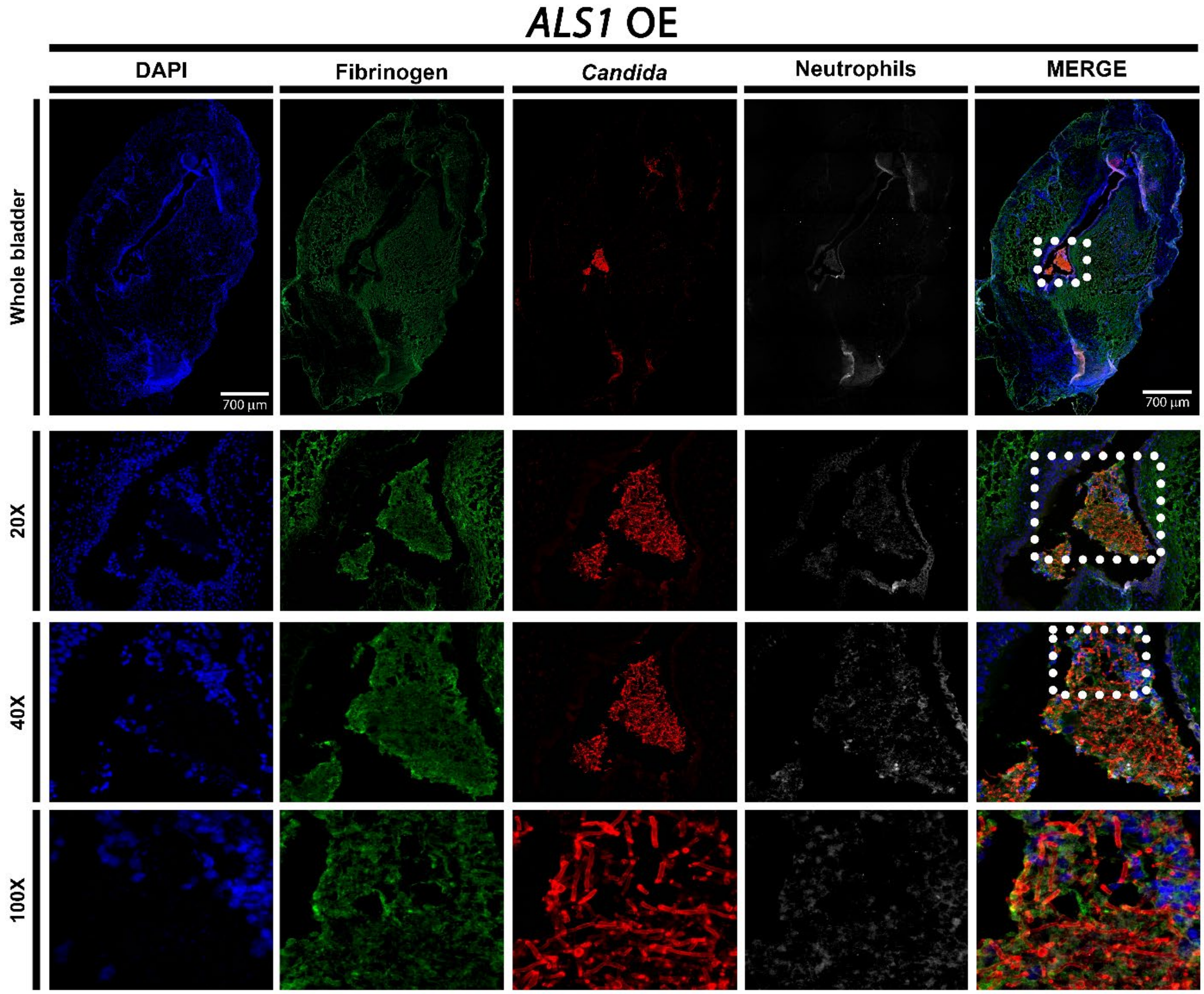


**Supplementary Figure 19.** ***C. albicans*** **SC5314** ***ALS1*** **OE bladder colonization during CAUTI.** Mice were implanted and infected with 1x $10^6$ CFUs with strain SC5314 *ALS1* OE. At 24 hpi, bladder tissues were harvested, fixed, and parafilm-embedded. Bladders were subjected to IF analysis using antibodies to detect Fg (anti-Fg; green), *C. albicans* (anti-*Candida*; red), and neutrophils (anti-Ly6G; white). Staining with DAPI (blue) delineated the urothelium and cell nuclei (representative images). IF stained bladder scale bars, 700 µm. White squares represent zoomed-in areas of higher magnification (20x, 40x, and 100x).

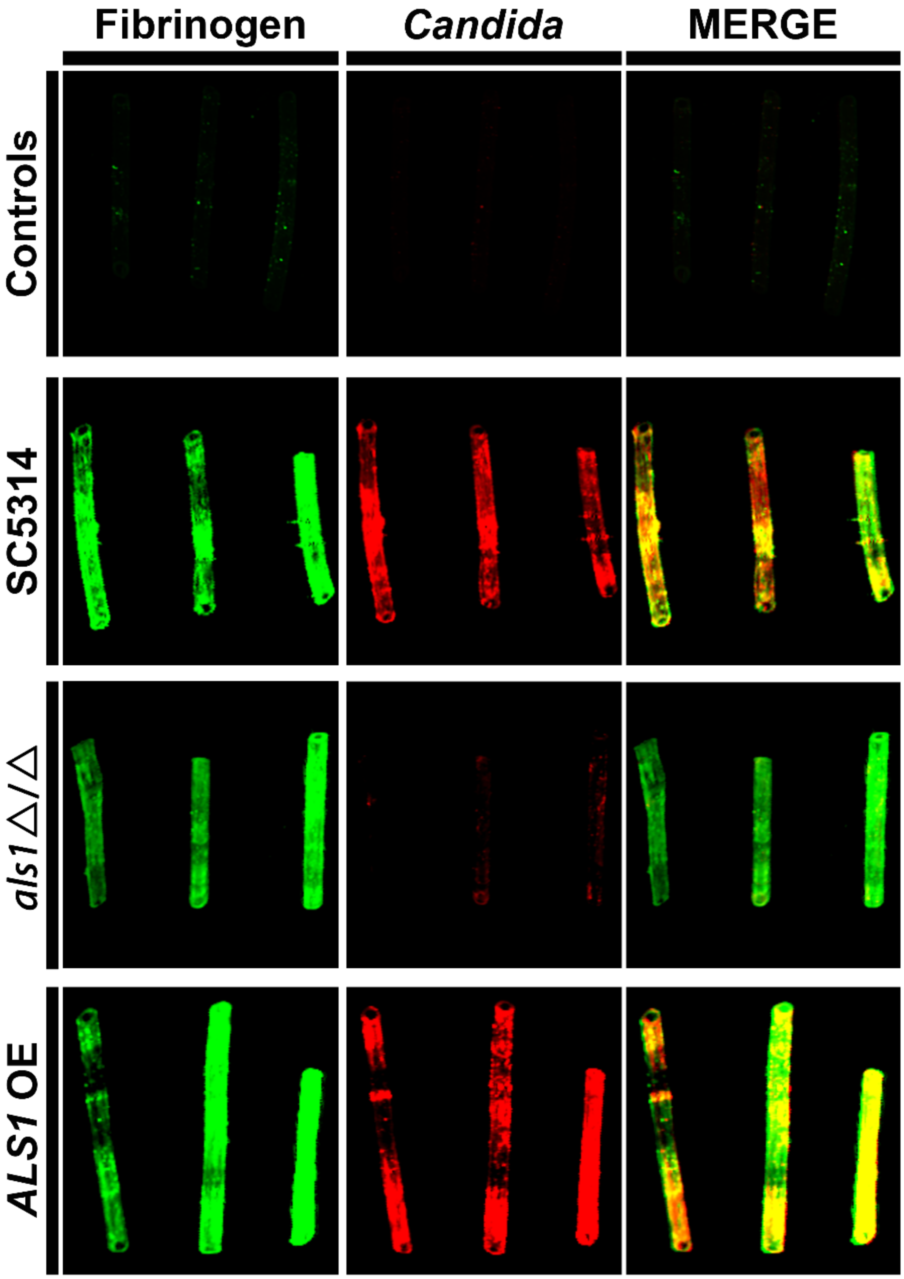


**Supplementary Figure 20. Colocalization of *C. albicans* strains with deposited fibrinogen on catheters during CAUTI.** Catheterized mice were challenged with 1x10$^6$ CFUs of the indicated *C. albicans* strain. Implanted catheters were retrieved 24 hpi and stained with antibodies to detect Fg (anti-Fg; green) and *C. albicans* (anti-*Candida*; red). Non-implanted catheters were used as negative controls (**A**).

**Supplementary Table 1.** Strains used in this study.

| Species | Strain | Description | Reference |
|---|---|---|---|
| *Candida albicans* | Pt62 | Clinical strain | (*1*) |
| *Candida albicans* | Pt65 | Clinical strain | (*2*) |
| *Candida albicans* | PNCL1 | Clinical strain | This study |
| *Candida albicans* | DAY286 | Laboratory strain | (*3*) |
| *Candida albicans* | SC5314 | Laboratory strain | (*4*) |
| *Candida albicans* | SC5314 *efg1*Δ/Δ*cph1*Δ/Δ | Laboratory strain | (*5*) |
| *Candida albicans* | SN425 –SC5314 background- | Laboratory strain | (*6*) |
| *Candida albicans* | *efg1*Δ/Δ (CJN2302) | Laboratory strain from Clarissa J. Nobile | (*6*) |
| *Candida albicans* | *EFG1 complement* (CJN2318) | Laboratory strain from Clarissa J. Nobile | (*6*) |
| *Candida albicans* | SN250- SC5314 background- Noble Library | Laboratory strain from Suzanne Noble Deletion Library | (*7*) |
| *Candida albicans* | *cph1*Δ/Δ | Laboratory strain from Suzanne Noble Deletion Library | (*7*) |
| *Candida albicans* | SC5314 WT adhesin library | Laboratory strain from Rebecca Shapiro Adhesin Deletion Library | (*8*) |
| *Candida albicans* | *als1*Δ/Δ | Laboratory strain from Rebecca Shapiro Adhesin Deletion Library | (*8*) |
| *Candida albicans* | *als3*Δ/Δ | Laboratory strain from Rebecca Shapiro Adhesin Deletion Library | (*8*) |
| *Candida albicans* | *als5*Δ/Δ | Laboratory strain from Rebecca Shapiro Adhesin Deletion Library | (*8*) |

| | | | |
|---|---|---|---|
| *Candida albicans* | *als7*Δ/Δ | Laboratory strain from Rebecca Shapiro Adhesin Deletion Library | (*8*) |
| *Candida albicans* | *als9*Δ/Δ | Laboratory strain from Rebecca Shapiro Adhesin Deletion Library | (*8*) |
| *Candida albicans* | *eap1*Δ/Δ | Laboratory strain from Rebecca Shapiro Adhesin Deletion Library | (*8*) |
| *Candida albicans* | *hwp1*Δ/Δ | Laboratory strain from Rebecca Shapiro Adhesin Deletion Library | (*8*) |
| *Candida albicans* | *hwp2*Δ/Δ | Laboratory strain from Rebecca Shapiro Adhesin Deletion Library | (*8*) |
| *Candida albicans* | *hyr1*Δ/Δ | Laboratory strain from Rebecca Shapiro Adhesin Deletion Library | (*8*) |
| *Candida albicans* | *iff4*Δ/Δ | Laboratory strain from Rebecca Shapiro Adhesin Deletion Library | (*8*) |
| *Candida albicans* | *rbr1*Δ/Δ | Laboratory strain from Rebecca Shapiro Adhesin Deletion Library | (*8*) |
| *Candida albicans* | *rbt1*Δ/Δ | Laboratory strain from Rebecca Shapiro Adhesin Deletion Library | (*8*) |
| *Candida albicans* | *als1*Δ/Δ (CJN2841) | Laboratory strain from Clarissa J. Nobile | This study |
| *Candida albicans* | *ALS1* complement (CJN2842) | Laboratory strain from Clarissa J. Nobile | This study |
| *Candida albicans* | SC5314 *ALS1* Overexpression | Laboratory strain from Rebecca Shapiro | (*9*) |

| | | | |
|---|---|---|---|
| *Candida albicans* | SC5314 CRISPR Overexpression Control | Laboratory strain from Rebecca Shapiro | (*9*) |

**Supplementary Table 2.** Percentage of *C. albicans* cell type in YPD and urine.

| Strain | Condition | Hyphae (%) | Pseudohyphae (%) | Yeast (%) |
|---|---|---|---|---|
| Pt62 | YPD 0h | 0 ± 0 | 0.4 ± 1.3 | 99.6 ± 1.3 |
| | YPD 24h | 0 ± 0 | 1.5 ± 3.2 | 98.5 ± 3.2 |
| | YPD 48h | 0 ± 0 | 0.5 ± 0.9 | 99.5 ± 0.9 |
| | YPD + serum 0h | 0 ± 0 | 0.4 ± 1.3 | 99.6 ± 1.3 |
| | YPD + serum 24h | 6.7 ± 11.9 | 3.3 ± 3.7 | 89.9 ± 14.4 |
| | YPD + serum 48h | 11.2 ± 8.7 | 2.5 ± 3.4 | 86.2 ± 9.5 |
| | Urine 0h | 0 ± 0 | 0 ± 0 | 100 ± 0 |
| | Urine 24h | 5.4 ± 7.6 | 9.1 ± 6.3 | 85.5 ± 8.7 |
| | Urine 48h | 0.4 ± 1.1 | 10.8 ± 7.2 | 88.9 ± 7.2 |
| | Urine + serum 0h | 0 ± 0 | 0.4 ± 1.3 | 99.6 ± 1.3 |
| | Urine + serum 24h | 25.6 ± 34.3 | 13.9 ± 11.2 | 60.5 ± 31.8 |
| | Urine + serum 48h | 22.5 ± 21.6 | 7.7 ± 6.3 | 69.8 ± 20.0 |
| Pt65 | YPD 0h | 0 ± 0 | 0 ± 0 | 100 ± 0 |
| | YPD 24h | 0 ± 0 | 0 ± 0 | 100 ± 0 |
| | YPD 48h | 0 ± 0 | 0.9 ± 2.6 | 99.1 ± 2.6 |
| | YPD + serum 0h | 0 ± 0 | 0 ± 0 | 100 ± 0 |
| | YPD + serum 24h | 8.5 ± 15.2 | 6.0 ± 11.1 | 85.5 ± 19.5 |
| | YPD + serum 48h | 2.3 ± 4.2 | 3.7 ± 4.7 | 93.9 ± 7.8 |
| | Urine 0h | 0 ± 0 | 0 ± 0 | 100 ± 0 |
| | Urine 24h | 0.7 ± 2.2 | 10.2 ± 7.7 | 89.0 ± 0 |
| | Urine 48h | 4.9 ± 10.9 | 10.1 ± 11.3 | 85.0 ± 11.7 |
| | Urine + serum 0h | 0 ± 0 | 0 ± 0 | 100 ± 0 |
| | Urine + serum 24h | 13.8 ± 20.6 | 13.9 ± 6.8 | 72.2 ± 24.2 |
| | Urine + serum 48h | 16.0 ± 31.1 | 12.3 ± 14.6 | 71.7 ± 34.2 |
| PCNL1 | YPD 0h | 0 ± 0 | 0 ± 0 | 100 ± 0 |
| | YPD 24h | 0 ± 0 | 3.0 ± 6.6 | 97.0 ± 6.6 |
| | YPD 48h | 0 ± 0 | 0.3 ± 1.0 | 99.7 ± 1.0 |
| | YPD + serum 0h | 0 ± 0 | 0 ± 0 | 100 ± 0 |
| | YPD + serum 24h | 6.0 ± 5.9 | 3.9 ± 3.1 | 90.2 ± 7.5 |
| | YPD + serum 48h | 9.0 ± 11.2 | 5.4 ± 8.7 | 85.6 ± 18.4 |
| | Urine 0h | 0 ± 0 | 0 ± 0 | 100 ± 0 |
| | Urine 24h | 2.5 ± 5.2 | 17.5 ± 13.3 | 80.0 ± 14.2 |

| | | | | |
|---|---|---|---|---|
| | Urine 48h | 5.1 ± 11.9 | 11.9 ± 9.8 | 83.0 ± 16.1 |
| | Urine + serum 0h | 0 ± 0 | 1.6 ± 4.8 | 98.4 ± 4.8 |
| | Urine + serum 24h | 25.8 ± 37.0 | 19.7 ± 18.7 | 54.6 ± 31.3 |
| | Urine + serum 48h | 18.0 ± 25.3 | 9.3 ± 10.7 | 72.8 ± 29.5 |
| DAY286 | YPD 0h | 0 ± 0 | 1.2 ± 3.7 | 98.8 ± 3.7 |
| | YPD 24h | 2.5 ± 5.6 | 3.2 ± 3.8 | 94.4 ± 7.2 |
| | YPD 48h | 1.1 ± 2.5 | 4.5 ± 5.3 | 94.4 ± 7.4 |
| | YPD + serum 0h | 1.0 ± 3.0 | 1.6 ± 3.5 | 97.4 ± 4.2 |
| | YPD + serum 24h | 15.9 ± 12.2 | 13.6 ± 6.9 | 70.5 ± 14.7 |
| | YPD + serum 48h | 20.4 ± 6.9 | 13.9 ± 4.8 | 65.7 ± 9.8 |
| | Urine 0h | 0 ± 0 | 2.7 ± 4.6 | 97.3 ± 4.6 |
| | Urine 24h | 22.2 ± 25.8 | 21.1 ± 27.6 | 56.6 ± 31.3 |
| | Urine 48h | 34.6 ± 19.1 | 18.0 ± 16.1 | 47.4 ± 27.1 |
| | Urine + serum 0h | 1.0 ± 3.0 | 0 ± 0 | 99.0 ± 3.0 |
| | Urine + serum 24h | 45.9 ± 31.1 | 21.9 ± 18.8 | 32.2 ± 19.5 |
| | Urine + serum 48h | 39.9 ± 27.7 | 13.6 ± 10.5 | 46.5 ± 23.5 |
| SC5314 | YPD 0h | 1.4 ± 4.2 | 1.6 ± 4.8 | 97.0 ± 5.9 |
| | YPD 24h | 3.8 ± 7.9 | 2.6 ± 3.7 | 93.6 ± 11.4 |
| | YPD 48h | 2.8 ± 5.9 | 2.1 ± 3.8 | 95.1 ± 7.8 |
| | YPD + serum 0h | 0 ± 0 | 1.6 ± 4.8 | 98.4 ± 4.8 |
| | YPD + serum 24h | 17.2 ± 13.4 | 11.4 ± 8.7 | 71.4 ± 11.8 |
| | YPD + serum 48h | 20.6 ± 23.1 | 9.0 ± 3.6 | 70.4 ± 25.1 |
| | Urine 0h | 1.4 ± 4.2 | 0 ± 0 | 98.6 ± 4.2 |
| | Urine 24h | 46.2 ± 26.2 | 18.2 ± 22.7 | 35.5 ± 9.7 |
| | Urine 48h | 36.8 ± 25.9 | 10.4 ± 10.1 | 52.8 ± 30.2 |
| | Urine + serum 0h | 1.4 ± 4.2 | 1.6 ± 4.8 | 97.0 ± 5.9 |
| | Urine + serum 24h | 52.2 ± 19.3 | 26.6 ± 14.3 | 21.2 ± 15.3 |
| | Urine + serum 48h | 38.1 ± 33.0 | 22.0 ± 18.7 | 39.9 ± 25.7 |
| SC5314 *efg1*Δ/Δ *cph1*Δ/Δ | YPD 0h | 0 ± 0 | 2.2 ± 3.2 | 97.8 ± 3.2 |
| | YPD 24h | 0 ± 0 | 1.0 ± 1.3 | 99.0 ± 1.3 |
| | YPD 48h | 0 ± 0 | 1.7 ± 3.3 | 98.3 ± 3.3 |
| | YPD + serum 0h | 0 ± 0 | 0.6 ± 0.9 | 99.4 ± 0.9 |
| | YPD + serum 24h | 0 ± 0 | 1.1 ± 2.7 | 98.9 ± 2.7 |

| | | | | |
|---|---|---|---|---|
| | YPD + serum 48h | 0 ± 0 | 0.4 ± 0.9 | 99.6 ± 0.9 |
| | Urine 0h | 0 ± 0 | 0.6 ± 0.9 | 99.4 ± 0.9 |
| | Urine 24h | 0 ± 0 | 15.9 ± 30.6 | 84.1 ± 30.1 |
| | Urine 48h | 0 ± 0 | 0.8 ± 1.3 | 99.2 ± 1.3 |
| | Urine + serum 0h | 0 ± 0 | 2.2 ± 3.0 | 97.8 ± 3.0 |
| | Urine + serum 24h | 0 ± 0 | 10.3 ± 16.8 | 89.7 ± 16.8 |
| | Urine + serum 48h | 0 ± 0 | 2.3 ± 6.3 | 97.7 ± 6.3 |
| SN250 (SC5314 background) Homann library | YPD 0h | 0.3 ± 0.7 | 0.8 ± 1.0 | 98.9 ± 1.5 |
| | YPD 24h | 0 ± 0 | 0.7 ± 1.4 | 99.3 ± 1.4 |
| | YPD 48h | 0.1 ± 0.2 | 0.6 ± 0.6 | 99.3 ± 0.7 |
| | YPD + serum 0h | 0.1 ± 0.4 | 1.0 ± 1.7 | 98.9 ± 1.7 |
| | YPD + serum 24h | 3.5 ± 2.6 | 3.7 ± 3.1 | 92.8 ± 4.8 |
| | YPD + serum 48h | 2.9 ± 4.2 | 4.2 ± 2.4 | 93.0 ± 5.2 |
| | Urine 0h | 0.8 ± 1.2 | 1.6 ± 1.7 | 97.6 ± 2.8 |
| | Urine 24h | 0.8 ± 2.3 | 3.2 ± 3.2 | 96.0 ± 4.1 |
| | Urine 48h | 4.2 ± 4.4 | 1.7 ± 1.9 | 94.0 ± 5.1 |
| | Urine + serum 0h | 0.1 ± 0.2 | 1.1 ± 1.6 | 98.9 ± 1.7 |
| | Urine + serum 24h | 2.5 ± 2.9 | 2.8 ± 6.6 | 94.6 ± 6.5 |
| | Urine + serum 48h | 30.1 ± 26.4 | 5.2 ± 6.7 | 64.6 ± 23.1 |
| *efg1*Δ/Δ | YPD 0h | 0 ± 0 | 0 ± 0 | 100 ± 0 |
| | YPD 24h | 0 ± 0 | 0 ± 0 | 100 ± 0 |
| | YPD 48h | 0 ± 0 | 0 ± 0 | 100 ± 0 |
| | YPD + serum 0h | 0 ± 0 | 0 ± 0 | 100 ± 0 |
| | YPD + serum 24h | 0 ± 0 | 0 ± 0 | 100 ± 0 |
| | YPD + serum 48h | 0 ± 0 | 0 ± 0 | 99.9 ± 0.1 |
| | Urine 0h | 0 ± 0 | 0.2 ± 0.3 | 99.8 ± 0.3 |
| | Urine 24h | 0 ± 0 | 0 ± 0 | 100 ± 0 |
| | Urine 48h | 0 ± 0 | 0 ± 0 | 100 ± 0 |
| | Urine + serum 0h | 0 ± 0 | 0.4 ± 0.8 | 99.6 ± 0.8 |
| | Urine + serum 24h | 0 ± 0 | 1.5 ± 4.1 | 98.5 ± 4.1 |
| | Urine + serum 48h | 0 ± 0 | 0.6 ± 1.7 | 99.4 ± 1.7 |
| SN250 (SC5314 background) | YPD 0h | 0 ± 0.1 | 0.3 ± 0.3 | 99.7 ± 0.4 |
| | YPD 24h | 0 ± 0 | 0 ± 0 | 100 ± 0 |
| | YPD 48h | 0 ± 0 | 0 ± 0 | 100 ± 0 |

| | | | | |
|---|---|---|---|---|
| Noble library | YPD + serum 0h | 0 ± 0 | 0.4 ± 0.6 | 99.6 ± 0.6 |
| | YPD + serum 24h | 1.6 ± 1.7 | 3.6 ± 3.2 | 94.8 ± 3.1 |
| | YPD + serum 48h | 2.7 ± 3.1 | 1.8 ± 1.7 | 95.5 ± 4.5 |
| | Urine 0h | 0 ± 0 | 0.1 ± 0.2 | 99.9 ± 0.2 |
| | Urine 24h | 2.6 ± 3.5 | 1.9 ± 1.8 | 95.6 ± 4.1 |
| | Urine 48h | 4.4 ± 3.9 | 2.5 ± 3.0 | 93.1 ± 4.8 |
| | Urine + serum 0h | 0 ± 0 | 0.2 ± 0.5 | 99.8 ± 0.5 |
| | Urine + serum 24h | 14.6 ± 30.3 | 6.0 ± 6.7 | 79.4 ± 28.1 |
| | Urine + serum 48h | 11.0 ± 13.8 | 5.6 ± 9.5 | 83.4 ± 19.0 |
| *cph1*Δ/Δ | YPD 0h | 0 ± 0 | 0 ± 0 | 100 ± 0 |
| | YPD 24h | 3.3 ± 7.7 | 4.1 ± 4.9 | 92.7 ± 8.6 |
| | YPD 48h | 2.0 ± 2.6 | 4.6 ± 7.9 | 93.4 ± 7.4 |
| | YPD + serum 0h | 0 ± 0 | 0.4 ± 0.8 | 99.6 ± 0.8 |
| | YPD + serum 24h | 35.2 ± 38.5 | 9.7 ± 9.9 | 55.1 ± 35.0 |
| | YPD + serum 48h | 9.5 ± 9.3 | 10.1 ± 9.1 | 80.4 ± 16.8 |
| | Urine 0h | 0 ± 0 | 0 ± 0 | 100 ± 0 |
| | Urine 24h | 23.9 ± 35.7 | 7.8 ± 17.2 | 68.3 ± 38.1 |
| | Urine 48h | 28.2 ± 28.0 | 9.8 ± 14.2 | 61.9 ± 24.0 |
| | Urine + serum 0h | 0 ± 0 | 0.2 ± 0.7 | 99.8 ± 0.7 |
| | Urine + serum 24h | 28.5 ± 43.0 | 29.6 ± 45.5 | 41.8 ± 44.9 |
| | Urine + serum 48h | 33.6 ± 42.4 | 7.1 ± 18.4 | 59.3 ± 47.2 |
| SC5314 WT Adhesin library | YPD 0h | 0.3 ± 0.9 | 2.3 ± 1.8 | 96.3 ± 2.1 |
| | YPD 24h | 1.7 ± 2.7 | 2.0 ± 3.3 | 96.3 ± 5.7 |
| | YPD 48h | 3.3 ± 5.2 | 4.6 ± 1.6 | 93.2 ± 6.2 |
| | YPD + serum 0h | 0.3 ± 0.6 | 2.4 ± 2.1 | 97.4 ± 2.1 |
| | YPD + serum 24h | 1.1 ± 1.5 | 5.1 ± 4.8 | 93.8 ± 5.5 |
| | YPD + serum 48h | 2.0 ± 3.2 | 7.3 ± 4.4 | 90.6 ± 5.0 |
| | Urine 0h | 0 ± 0 | 2.1 ± 1.6 | 97.9 ± 1.6 |
| | Urine 24h | 14.0 ± 8.0 | 3.3 ± 4.1 | 82.7 ± 9.8 |
| | Urine 48h | 13.4 ± 6.7 | 1.0 ± 2.5 | 85.6 ± 5.6 |
| | Urine + serum 0h | 0.9 ± 1.3 | 3.3 ± 1.9 | 95.8 ± 2.5 |
| | Urine + serum 24h | 19.6 ± 21.0 | 0.3 ± 1.0 | 80.0 ± 20.9 |

| | | | | |
|---|---|---|---|---|
| | Urine + serum 48h | 26.9 ± 14.8 | 3.1 ± 4.3 | 70.0 ± 15.1 |
| *als1*Δ/Δ | YPD 0h | 0.2 ± 0.4 | 4.6 ± 3.9 | 95.2 ± 3.8 |
| | YPD 24h | 1.0 ± 3.0 | 5.6 ± 3.5 | 93.4 ± 5.0 |
| | YPD 48h | 1.5 ± 2.9 | 17.6 ± 15.6 | 80.9 ± 15.5 |
| | YPD + serum 0h | 0.4 ± 0.8 | 5.7 ± 4.2 | 93.9 ± 4.2 |
| | YPD + serum 24h | 4.7 ± 7.0 | 22.0 ± 8.2 | 73.2 ± 12.9 |
| | YPD + serum 48h | 4.1 ± 4.4 | 20.1 ± 9.2 | 75.8 ± 11.2 |
| | Urine 0h | 0.1 ± 0.2 | 4.0 ± 3.0 | 95.9 ± 2.9 |
| | Urine 24h | 2.7 ± 4.4 | 13.6 ± 13.0 | 83.6 ± 11.9 |
| | Urine 48h | 4.0 ± 6.1 | 18.9 ± 10.1 | 77.1 ± 9.5 |
| | Urine + serum 0h | 0.2 ± 0.3 | 2.9 ± 1.0 | 96.9 ± 0.9 |
| | Urine + serum 24h | 9.4 ± 17.3 | 11.4 ± 12.7 | 79.2 ± 25.3 |
| | Urine + serum 48h | 22.0 ± 32.1 | 9.5 ± 9.8 | 68.5 ± 27.7 |

**Supplementary Table 3.** Percentage of catheter colonization by *C. albicans* strains during CAUTI.

| *C. albicans* strain | Catheter colonization (%) |
|---|---|
| Pt62 | 59.2[a] ± 17.7[b] |
| Pt65 | 60.8 ± 18.2 |
| PCNL1 | 72.7 ± 16.5 |
| DAY286 | 74.9 ± 15.6 |
| SC3514 WT | 78.9 ± 14.0 |
| SC5314 *efg1*Δ/Δ*cph1*Δ/Δ | 10.4 ± 7.5 |
| SC5314 WT (Adhesin library) | 92.5 ± 3.7 |
| *als1*Δ/Δ | 2.8 ± 2.4 |
| *ALS1* OE | 87.5 ± 10.3 |

[a] mean; [b] standard deviation.

**Supplementary Table 4.** Primers used in this study

| GENE | Forward primer (5'-3') | Reverse primer (5'-3') |
|---|---|---|
| EFG1 | GGTCAGTATAATGCTCCTGGTAAG | CAGCACCACCCTGGTAATAAT |
| ALS1 | GACTAGTGAACCAACAAATACCAG | ACCAGAAGAAACAGCAGGTG |
| ITS2 | TGGGTTTGCTTGAAAGACGC | CCGCCGCAAGCAATGTTTTT |

.

**Supplementary Table 5.** Flow antibodies used to quantify neutrophil populations.

| Molecule | Clone | Fluorophore | Vendor |
|---|---|---|---|
| CD45 | 30-F11 | PE-Cy5 | BD Bioscience |
| CD11b | M1/70 | BV650 | BioLegend |
| Ly6G | 1A8 | BV711 | BD Bioscience |
| Live/Dead | | APC | Thermo Scientific |
